# Transcript errors generate a continuous stream of amyloid and prion-like proteins in human cells

**DOI:** 10.1101/2023.05.11.540433

**Authors:** Claire S. Chung, Yi Kou, Sarah J. Shemtov, Bert M. Verheijen, Ilse Flores, Kayla Love, Ashley Del Dosso, Max Thorwald, Yuchen Liu, Renaldo G. Toney, Lucy Carrillo, Megan Nguyen, Huang Biao, Yuxin Jin, Ashley Michelle Jauregui, Juan Diaz Quiroz, Darcie L. Moore, Stephen Simpson, Kelley Thomas, Marcelo P. Coba, Zhongwei Li, Bérénice A. Benayoun, Joshua Rosenthal, Scott Kennedy, Giorgia Quadrato, Jean-Francois Gout, Lin Chen, Marc Vermulst

## Abstract

Aging is characterized by the accumulation of amyloid and prion-like proteins. However, the molecular mechanisms by which these proteins arise remain unclear. Here, we demonstrate that transcript errors generate amyloid and prion-like proteins in a wide variety of human cell types, including stem cells, brain organoids, and fully differentiated neurons. Intriguingly, some of these proteins are identical to proteins previously implicated in familial cases of amyloid diseases, raising the possibility that both familial and non-familial cases are caused by identical mutant proteins. However, transcript errors also generate amyloid proteins that have not been observed before, suggesting that aging cells are exposed to a second class of pathogenic proteins we are currently unaware of. Finally, we show that transcript errors are readily generated by DNA damage, a hallmark of human aging and a staple of multiple proteotoxic diseases, including Alzheimer’s disease. Together, these observations greatly expand our understanding of mutagenesis in human aging and disease and suggest a new mechanism by which amyloid diseases can develop.

## INTRODUCTION

Protein aggregation is a defining hallmark of human aging and disease(*1, 2*). At a molecular level, protein aggregates are formed by misfolded proteins that form amorphous protein deposits or self-assemble into large, neatly organized amyloid fibers. These aggregates play an important role in various neurodegenerative diseases, including Alzheimer’s disease (AD), Parkinson’s disease (PD) and Creutzfeld-Jakob Disease (CJD)(*3, 4*). However, they also contribute to the functional decline associated with normal aging and the pathology of a wide variety of other age-related diseases, including cancer(*5*), amyotrophic lateral sclerosis, diabetes, heart disease and cataracts(*6–9*). In familial cases of amyloid diseases, patients tend to carry a single point mutation that dramatically increases the amyloid propensity of the affected protein(*10*). However, why proteins misfold and aggregate in non-familial cases of amyloid diseases remains unclear.

One long-standing hypothesis is that in non-familial cases of these diseases, amyloid proteins are generated by epi-mutations, non-genetic mutations that are only present in transcripts. For example, if a mistake was made during RNA synthesis(*11–14*) or RNA editing(*15*), a small cache of mutant proteins would be generated that could display amyloid or prion-like behavior. Although their initial number would be small, amyloid and prion-like proteins are defined by their ability to replicate themselves by binding to WT proteins through strong, non-covalent interactions and converting them to an amyloid state(*16*). Through this self-templating mechanism, a small cache of mutant proteins could rapidly grow in size and number, and eventually seed the amyloid fibers that characterize aging cells (**fig. 1a**).

**Figure 1.**
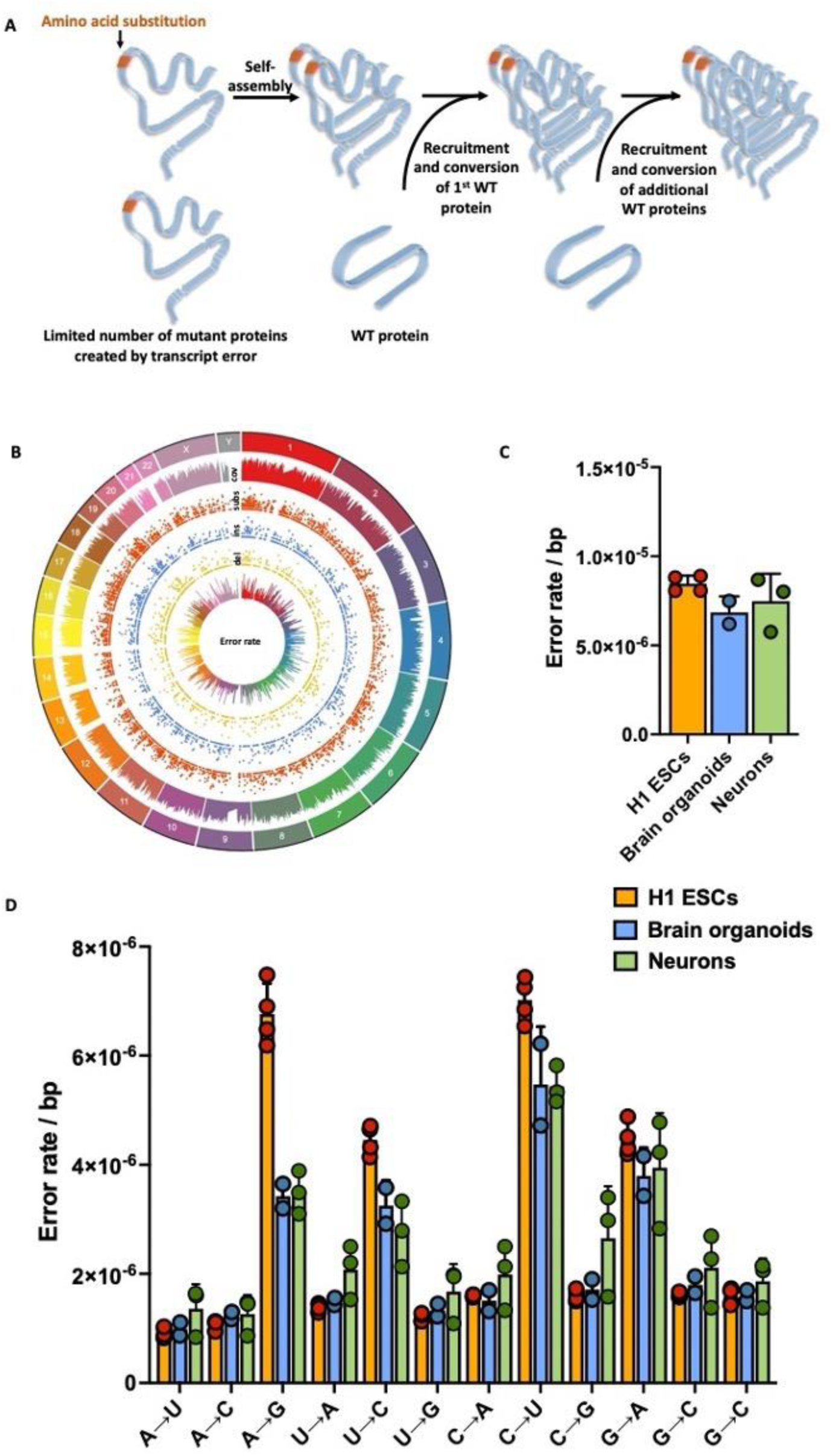
Graphical representation of hypothesis and summary of transcription error data. **A.** We hypothesize that transcription errors could give rise to amyloid and prion-like proteins. This relatively small cache of mutant proteins can then form an amyloid seed that recruits WT proteins and converts them to an amyloid state to generate large amyloid fibers. **B.** Transcription errors were identified across the genome of H1 Human embryonic stem cells, brain organoids and human neurons. **C-D.** The error rate and spectrum of H1 Human embryonic stem cells, brain organoids and human neurons is nearly identical. Error bars indicate standard error of the mean.

However, transcript errors are exceedingly difficult to detect, which has made it difficult to test this hypothesis in a comprehensive manner. To solve this problem, we recently optimized a new RNA sequencing tool termed circle-sequencing(*17*), which allows for high-fidelity sequencing of mRNA molecules (**fig. S1**). Here, we use circle-sequencing to demonstrate that transcript errors are ubiquitous in human cells, and that they indeed result in proteins with amyloid and prion-like properties. We support these observations with a variety of cellular, biochemical and biophysical experiments that demonstrate that the proteins generated by these errors can successfully convert WT proteins to an amyloid state, which then self-assemble into neatly organized amyloid fibers. Finally, we show that the amount of mutant proteins required to initiate large-scale protein aggregation is routinely breached as a result of DNA damage, a ubiquitous hallmark of aging cells. As a result, our experiments establish a direct, mechanistic link between DNA damage and protein aggregation, two of the major hallmarks of human aging and age-related diseases, including Alzheimer’s disease. In doing so, our experiments redefine the role of mutagenesis in human aging and disease, and suggest a new mechanism by which amyloid and prion diseases can develop.

## RESULTS

To test whether transcript errors give rise to amyloid or prion-like proteins, we probed the transcriptome of H1 human embryonic stem cells (H1 ESCs), brain organoids, neurons with circ-seq, a massively parallel sequencing approach that uses consensus sequencing to enable high-fidelity RNA sequencing(*17, 18*). The brain organoids and neurons we sequenced were generated directly from the H1 ESCs (**fig. S2**), so that the genetic background between these models was consistent and the results could be compared to each other. In addition, we sequenced the H1 ESCs at 300x coverage to generate a custom-made reference genome and ensure that single nucleotide polymorphisms or low-level mutations could be excluded from downstream analyses (**fig. S3**). In total, these sequencing efforts yielded >160,000 transcript errors that affected >11,000 genes across all three models (**fig. 1b**). A complete list of the errors we detected can be found in the supplemental material attached to this publication. Interestingly, each model displayed a relatively similar error rate and spectrum, suggesting that the error rate of transcription is relatively independent of cellular fate, proliferation rate and differentiation status (**fig. 1c**). We did observe a higher rate of A→G errors in the H1 ESCs cells though, which we previously found to reflect the impact of A to I RNA editing on the transcriptome(*19*).

We then used two approaches to determine if the errors we had detected result in amyloid or prion-like proteins: a literature-based approach and a bio-informatic approach. In our literature-based approach, we focused on 70 proteins that are directly implicated in various amyloid and prion-like diseases, including PRNP (CJD and Gerstman–Sträussler–Scheinker syndrome(*20*)), APP (AD)(*21*)), SOD1 and FUS (Amyotrophic Lateral Sclerosis(*22*), and TTR (transthyretin amyloidosis) (**table 1, table S1**). Over the past 3 decades, thousands of mutations have been identified in these proteins that cause familial cases of proteinopathies. In most cases, these mutations greatly increase the amyloid and prion-like potential of the affected proteins. We reasoned that if transcript errors generate identical mutant proteins, they are likely to generate amyloid and prion-like proteins as well. To test this idea, we cross-referenced the errors we detected with various databases that catalogue germline mutations implicated in amyloid diseases, including Clinvar(*23*) and the human genome mutation database(*24*). Of the 1936 errors that affected amyloid and prion-like proteins, we identified 38 errors that give rise to mutant proteins previously seen in the clinic. For example, 2 of the errors we detected generate mutant versions of the SOD1(*25*) and FUS protein(*26*), both of which were previously identified in familial cases of amyotrophic lateral sclerosis (ALS) (**table 1, table S1)**, while another error generated a mutant version of the human prion protein (PRNP^A133V^) that causes Gerstman–Sträussler–Scheinker syndrome (GSS)(*27*). Other errors generated pathological versions of TTR (amyloidogenic transthyretin amyloidosis), CSTB (progressive myoclonic epilepsy), TGFBI (corneal dystrophy), APP (AD), CRYGD (coralliform cataracts), TP53 (cancer), Medin (natural aging), and TUBA1A (tubulinopathies) and others. In addition, we identified 75 errors that affect key amino acids directly implicated in disease, although they mutated them to a different residue compared to the clinic. For example, one of these errors generates a mutant version of PRNP (PRNP^V210A^) that closely resembles a PRNP^V210I^ mutation known to be one of the most common causes of familial CJD(*28*) (both alanine and isoleucine are aliphatic amino acids). Similar errors were present in transcripts that encode APP, CSTB, HNRNPA1 (inclusion body myopathy with FTD), TGFBI, TP53, TTR and 10 other proteins (**table 1, table S1**). A substantial portion of these errors is likely to affect the amyloid and prion-like behavior of these proteins as well.

**Table 1.**
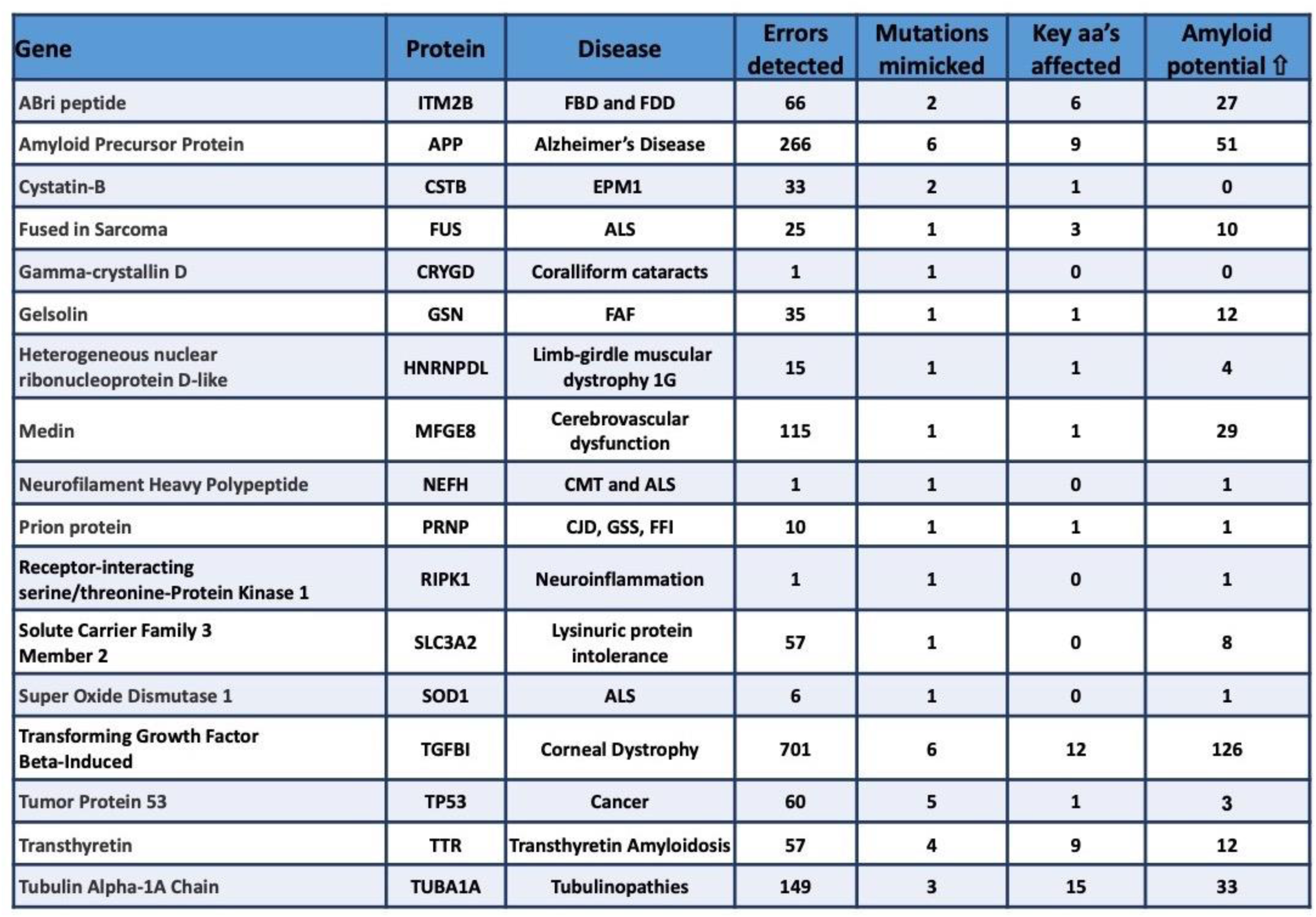
Transcription errors affect proteins directly implicated in amyloid and prion diseases. **Column 1:** Gene name. **Column 2:** Protein name. **Column 3:** Disease associated with protein. **Column 4:** Number of errors detected in transcripts that were derived from this gene. **Column 5:** Number of errors that generate mutant proteins identical to those seen in familial cases of amyloid diseases. **Column 6:** Number of errors that affect an amino acid (aa) known to be involved in disease, but mutate it to a different residue compared to the clinic. **Column 6:** Number of errors that increase the amyloid potential of these proteins as predicted by bio-informatic analysis (AmyPred-FRL).

To confirm that the errors we identified through our literature-based approach indeed result in proteins with amyloid behavior, we selected two candidates for follow-up experiments. One of these errors generates a mutant version of SOD1 (SOD1^G142E^, **fig. 2a-d**) while the second error generates a mutant version of FUS (FUS^R521H^, **fig. 2e-h**). These mutant proteins were previously identified in familial cases of ALS(*25, 26*). We expressed these proteins in primary human fibroblasts, HEK293 cells and glioblastoma cells by lentiviral transfection (**fig. 2, S4**) and then imaged them by confocal microscopy. Consistent with the idea that transcription errors generate mutant proteins that display amyloid behavior, we found that both SOD1^G142E^ and FUS^R521H^ aggregated in all 3 cell types, while the WT proteins did not. In addition, we found that the mutant SOD1 and FUS proteins were mislocalized. While WT FUS is predominantly present in the nucleus (where it aids RNA splicing, gene expression and DNA repair(*29*)), the mutant protein was completely excluded from the nucleus and formed large punctate deposits throughout the cytoplasm (**fig. 2e-h**). These observations complement similar results by others (*29–33*). Similarly, SOD1 is normally distributed throughout the cytoplasm and the nucleus, but we found that SOD1^G142E^ was excluded from the nucleus and formed large protein deposits in the cytoplasm (**fig. 2b-d**). Importantly, nuclear exclusion and protein aggregation of FUS and SOD1 are key components of the pathology associated with ALS(*34, 35*). Finally, we co-expressed WT and mutant SOD1 in the same cells and monitored their behavior. Intriguingly, we found that when co-expressed with SOD1^G142E^, WT SOD1 no longer distributed equally throughout the cells, but was excluded from the nucleus and assembled into the same amyloid deposits as SOD1^G142E^ (**fig. 2i-l**), strongly suggesting that WT SOD1 was recruited by SOD1^G142E^ and converted to an amyloid state. We made similar observations for WT and mutant FUS^R521H^ (**fig. 2l-o**). WT FUS was almost always excluded from the nucleus in the presence of FUS^R521H^, and sequestered in cytoplasmic deposits with FUS^R521H^, although rare exceptions did occur (**fig. S5**). Consistent with the idea that SOD1^G142E^ has amyloid properties, transmission electron microscopy (TEM) demonstrated that mutant SOD1 can form amyloid fibers *in vitro* (**fig. 2p, q**). When taken together, these experiments provide an important proof of principle for the idea that transcription errors give rise to amyloid proteins. Moreover, because RNAPII constantly generates new mRNA molecules inside cells, and transcription by RNAPII is relatively error prone, (the error rate of transcription is approximately >100-fold higher than the mutation rate(*36*)), we conclude that transcription errors generate a continuous stream of amyloid and prion-like proteins in human cells.

**Figure 2.**
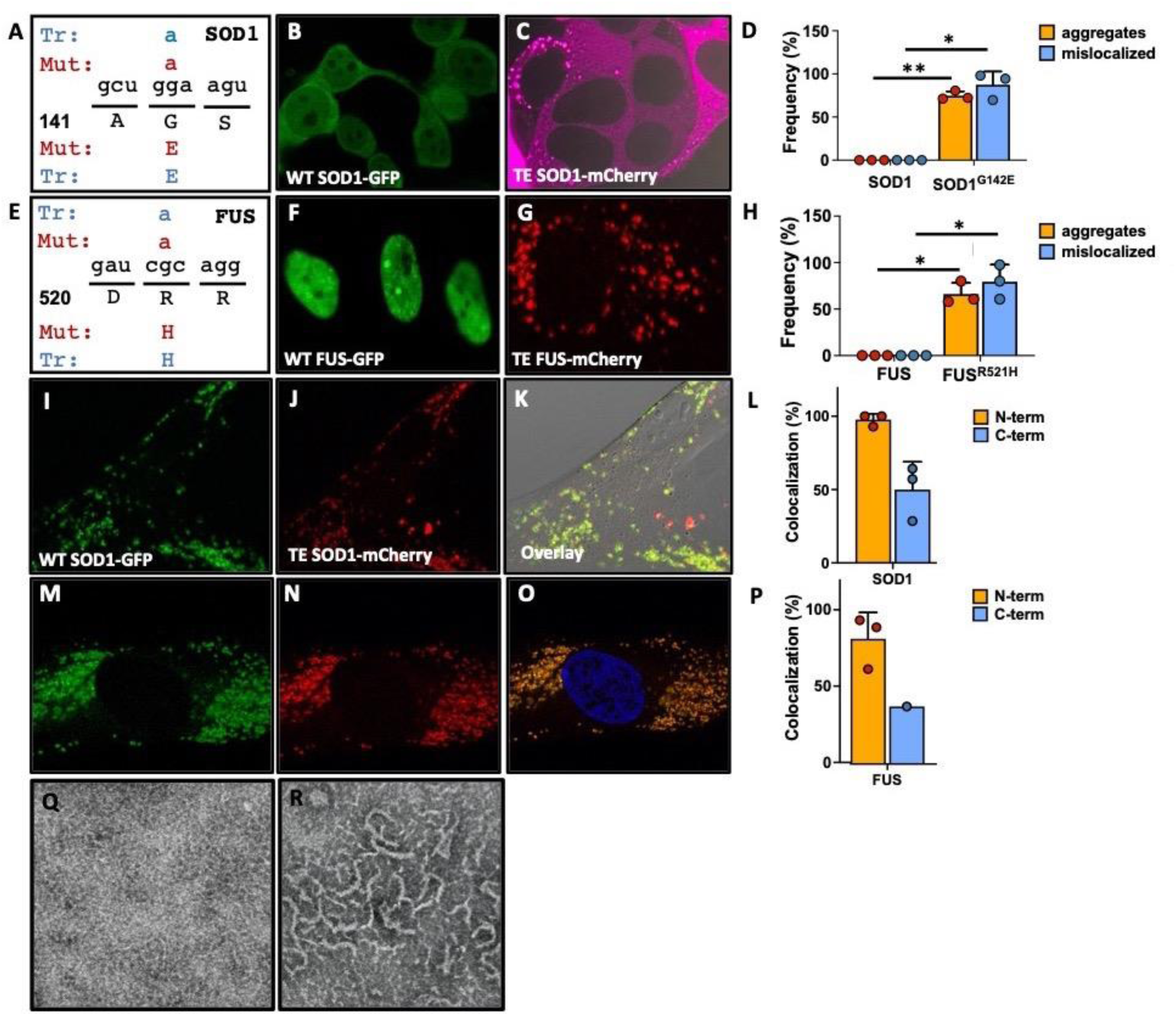
Transcription errors result in proteins with increased amyloid behavior. **A.** A transcription error (Tr) was identified in the SOD1 transcript that mimics a mutation (Mut) implicated in amyotrophic lateral sclerosis. This error substitutes a guanine for an adenine base, resulting in a glycine (G) to glutamine (E) mutation at residue 142. **B.** WT SOD1 is soluble and present throughout the cell, including the nucleus. **C.** In contrast, SOD1^G142E^ proteins form aggregates and are excluded the nucleus. **D.** Quantification of WT and mutant SOD1 aggregation and mislocalization. Depicted are the % of cells with aggregates or mislocalized proteins. **E.** A transcription error was identified in the FUS transcript that mimics a mutation implicated in amyotrophic lateral sclerosis. This error substituted a guanine for an adenine base, resulting in an arginine (R) to histidine (H) mutation at reside 521. **F.** WT FUS is present in a soluble state in the nucleus, while FUS^R521H^ (**G**) forms aggregates outside of the nucleus. **H.** Quantification of FUS aggregation and mislocalization. Depicted are the % of cells with aggregates or mislocalized proteins. **I-K**. When WT and SOD1^G142E^ are expressed simultaneously, WT SOD1 is excluded from the nucleus and recruited into extranuclear aggregates. **L.** Quantification of SOD1 colocalization with either N-terminal or C-terminal tags. **M-O.** WT and FUS^R521H^ co-expression in primary human fibroblasts, demonstrating that mutant and WT FUS co-localize in cytoplasmic aggregates. **P.** Quantification of FUS colocalization with either N-terminal or C-terminal tags. **Q.** WT SOD1 does not form fibers under TEM, but SOD1^G142E^ does (**R**). * =P<0.05. ** = P<0.01 according to an unpaired t-test with Welch’s correction. Error bars indicate standard error of the mean.

In addition to proteins directly connected to disease, we wondered whether transcription errors can also generate mutant proteins whose amyloid properties have not been characterized yet. To test this hypothesis, we used an unbiased bioinformatic approach to analyze the impact of errors on amyloid and prion-like proteins. First, we used AmyPred-FRL to analyze errors that affect amyloid proteins and found that 457 were predicted to increase their amyloid potential (**table 1, S1**). Second, we used the PAPA(*37*) to analyze errors that affect proteins with prion-like domains. Although the PRNP gene encodes the canonical prion protein in humans, many proteins are now known to contain prion-like domains. Mutations in these domains can greatly increase the prion-like behavior of these proteins in a wide variety of contexts, including proteotoxic diseases. For example, mutations in the prion-like domain of HNRNPA1 and HNRNPAB2 increase the amyloid behavior of these proteins and can cause multisystem proteinopathies and ALS(*38, 39*). By applying this algorithm to our dataset, we found that 393 transcript errors are predicted to display increased prion-like behavior (**table 2, table S2**).

**Table 2.** Transcription errors affect proteins with prion-like domains. **Column 1:** Gene name. **Column 2:** Protein name. **Column 3:** Location of prion-like domain inside protein. **Column 4:** Number of errors detected in transcripts that were derived from this gene. **Column 5:** Number of errors that affect the prion-like domain. **Column 6:** The number of errors that increase the prion-like potential of these proteins as predicted by bio-informatic analysis (PAPA).

| Gene | Protein | PrID | Errors detected | Errors in PrID | Prion potential ↑ |
| --- | --- | --- | --- | --- | --- |
| Annexin A11 | ANXA11 | 41-177 | 25 | 6 | 3 |
| Cell Cycle Associated Protein 1 | CAPRIN1 | 466-628 | 59 | 14 | 20 |
| DEAD-Box Helicase 5 | DDX5 | 531-614 | 64 | 3 | 14 |
| DEAD-Box Helicase 17 | DDX17 | 614-719 | 46 | 4 | 9 |
| Ewing Sarcoma breakpoint region 1 | EWSR1 | 1-256 | 68 | 21 | 17 |
| Fused in Sarcoma | FUS | 1-239 | 35 | 8 | 11 |
| Heterogeneous nuclear ribonucleoprotein A0 | HNRNPA0 | 206-305 | 23 | 7 | 3 |
| Heterogeneous nuclear ribonucleoprotein A1 | HNRNPA1 | 190-307 | 167 | 19 | 27 |
| Heterogeneous nuclear ribonucleoproteins A2/B1 | HNRNPA2B1 | 197-353 | 38 | 6 | 12 |
| Heterogeneous nuclear ribonucleoprotein A3 | HNRNPA3 | 207-378 | 36 | 6 | 14 |
| Heterogeneous nuclear ribonucleoprotein D0 | HNRNPD | 195-260 | 53 | 3 | 9 |
| Heterogeneous nuclear ribonucleoprotein H3 | HNRNPH3 | 268-346 | 15 | 3 | 4 |
| Heterogeneous nuclear ribonucleoprotein U | HNRNPU | 683-825 | 132 | 6 | 15 |
| Heterogeneous nuclear ribonucleoprotein U | HNRNPU | 683-825 | 132 | 6 | 15 |
| Heterogeneous nuclear ribonucleoprotein U-like 1 | HNRNPUL1 | 531-766 | 105 | 9 | 17 |
| Heterogeneous nuclear ribonucleoprotein U-like 2 | HNRNPUL2 | 641-747 | 21 | 4 | 5 |
| Interleukin enhancer-binding factor 3 | ILF3 | 661-894 | 67 | 7 | 19 |
| Nuclear receptor coactivator 1 | NCOA1 | 908-1198 | 8 | 3 | 5 |
| Nuclear factor of activated T-cells 5 | NFAT5 | 999-1454 | 12 | 6 | 4 |
| Nucleoporin 153 | NUP153 | 1322-1453 | 23 | 4 | 4 |
| Polyhomeotic Homolog 1 | PHC1 | 7-104 | 103 | 8 | 5 |
| R3H domain-containing protein 2 | R3HDM2 | 394-714 | 8 | 3 | 3 |
| SRY-Box Transcription Factor 2 | SOX2 | 154-231 | 14 | 4 | 0 |
| TAR DNA-binding protein 43 | TARDBP | 277-414 | 35 | 3 | 5 |
| TRK-fused gene protein | TFG | 223-400 | 24 | 8 | 6 |
| Yip1 domain family member 5; | YIPF5 | 1-85 | 12 | 4 | 1 |
| Zinc Finger MIZ-Type Containing 1 | ZMIZ1 | 248-403 | 24 | 4 | 5 |

Next, we extracted information from the Prionscan(*40*), PLAAC(*41*) and Amyloid Protein Database(*42*) to build a comprehensive database of proteins that have the potential to display amyloid and prion-like features. We then cross-referenced this database with the transcription errors we detected to identify errors that are likely to enhance these features. To test the veracity of these predictions, we examined errors that affect the TP53 protein in greater detail. TP53 is an essential tumor suppressor protein involved in DNA repair(34), transcription, cellular senescence and apoptosis, and aggregates in 15% of human cancers(5, 35–37). With the bio-informatic tools described above we identified 5 transcription errors that are likely to increase the amyloid propensity of TP53: TP53^S149F^, TP53^G245S^, TP53^G279A^, TP53^S315F^ and TP53^P318L^. When we mapped these mutations onto the crystal structure of TP53 we noticed that the S149F mutation (**fig. 3a**) is located in a loop at the edge of the TP53 β-sandwich core (loop 146-WVDSTPPPGTR-156). Based on its location and the structural change it introduces (**fig. 3b, c**), we predicted that this mutation may increase the amyloid propensity of the local peptide sequence (the 146-WVDSTPPPGTR-156 loop) and enhance the interaction between the β-sandwich cores of separate TP53 monomers, thereby leading to the assembly of the extended β-sheet structures that are characteristic of amyloid proteins (**fig. 3b, c**). Mutations in the loop at the edge of the β-sandwich core of the TTR protein were previously shown to promote amyloid formation through a similar structure-based mechanism(*43*). To test this hypothesis, we expressed the core domain (aa 92-292) of WT and mutant TP53 in bacterial cells and analyzed the behavior of these proteins by TEM. Consistent with our predictions, we found that TP53^S149F^ indeed aggregated into large protein deposits, while WT TP53 did not (**fig 3d, e**). These aggregates displayed Congo-red birefringence under polarized light, a strong indicator of amyloid formation (**fig. 3f, g**). We conclude that in addition to amyloid proteins directly implicated in disease, transcription errors also give rise to novel mutant proteins that tend to form amyloid structures.

**Figure 3.**
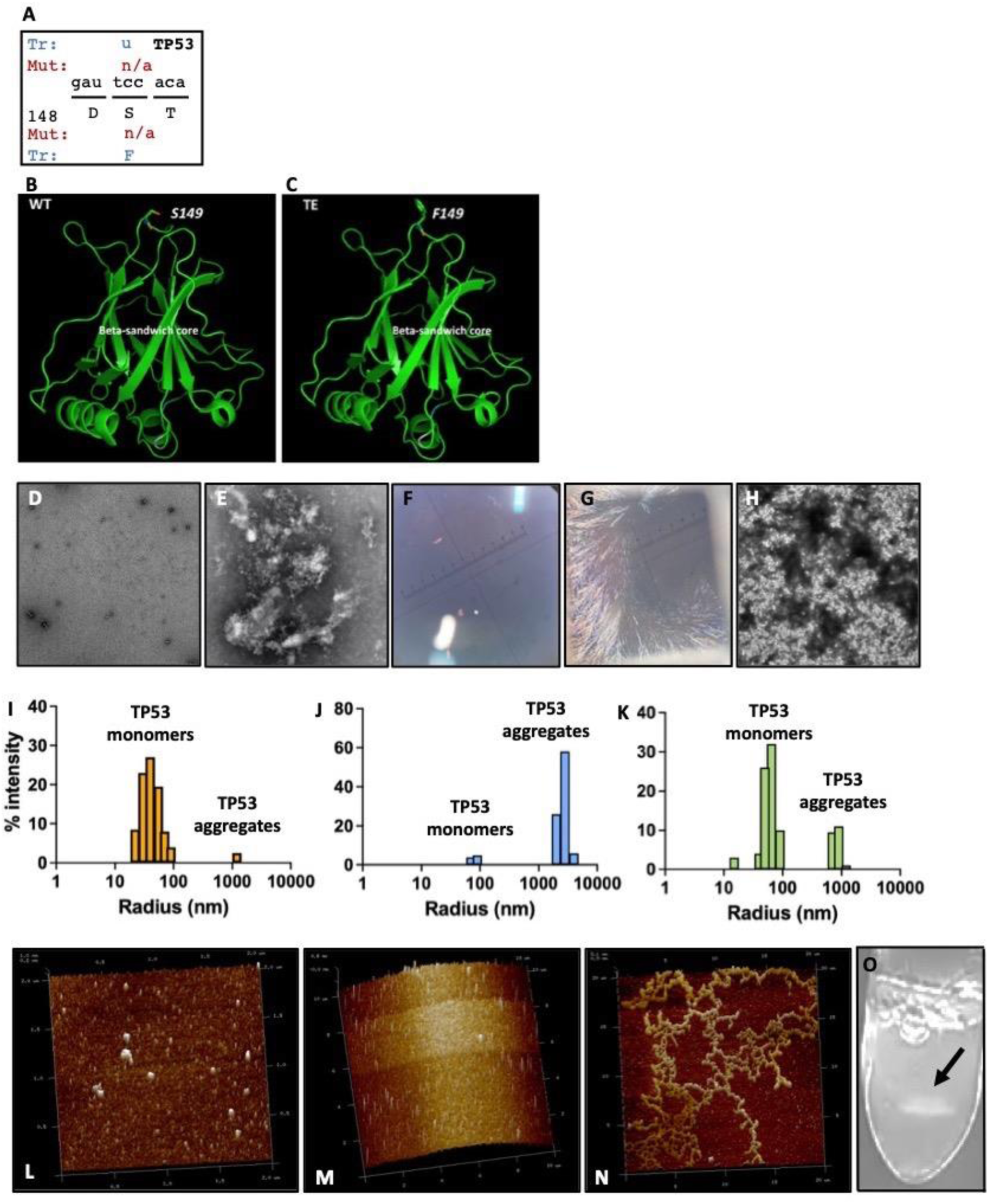
Biophysical examination of WT and mutant TP53. **A.** A transcription error (Tr) was identified in a TP53 transcript that substitutes a uracil for a cytosine base, resulting in a serine (S) to phenyl-alanine (F) mutation at residue 149. **B-C.** Predicted structure of WT (**B**) and mutant TP53 (**C**). **D.** Transmission electron microscopy showed little or no aggregates of WT TP53, while TP53^S149F^ induces large protein aggregates (**E**). **F-G**. Congo-red birefringence under polarized light indicates that TP53^S149F^ forms amyloid fibrils (**G**), while WT TP53 does not (**F**). **H.** After addition of 1% TP53^S149F^ to a solution of WT TP53 (v/v), the WT solution generated countless aggregates. **I-K.** Dynamic light scattering, which can be used determine the radius of protein particles, indicates that WT TP53 is primarily in a monomeric form (**I**), while mutant TP53 consists of aggregates greater than 1000nm (**J**). After 2% TP53^S149F^ is added to a solution of WT TP53 (v/v), a large amount of TP53 aggregates emerges (**K**). L. TP53^S149F^ aggregates were sonicated to create a seed solution of particles that are around 0.1µm in size, which equates to 800-1000 proteins. (**M**) WT TP53 solution shows no apparent aggregation; (**N**) Adding the TP53^S149F^ amyloid seed solution to WT TP53 in a 1:100 ratio induced fibril growth. **O.** Protein aggregates created by mutant TP53 form spontaneously and can be seen by the naked eye (arrow).

Next, we decided to test if TP53^S149F^ can convert WT TP53 to an amyloid state, similar to SOD1^G142E^ and FUS^R521H^, and if so, how much TP53^S149F^ was required to initiate this process. To answer this question, we added vanishing amounts of TP53^S149F^ to a WT TP53 solution and found by TEM that 1% of TP53^S149F^ (v/v) was sufficient to initiate the aggregation of the WT protein (**fig. 3h**). We confirmed these findings in a dynamic light scattering experiment (**fig. 3i-k**) that demonstrated that while WT TP53 was present at a size consistent with TP53 monomers, TP53^S149F^ aggregated into deposits that were >100-fold larger in size. Moreover, when we added 2% (v/v) TP53^S149F^ to the WT solution, we observed a disproportionate increase in TP53 aggregates that could only be explained by mutant-induced aggregation of WT proteins. Finally, we used atomic force microscopy (AFM) to characterize the cross-seeding behavior between WT and mutant TP53 further. First, we prepared a seeding solution of TP53^S149F^ aggregates by sonication and centrifugation with an average particle size of 0.1 µm as determined by multi-angle light scattering (MALS) and AFM. Particles that are 0.1 µm in size are roughly equivalent to ~800-1000 molecules, a number that could be generated by the translation of 1 or a few mutant transcripts (**fig. 3l**). We then mixed these particles into the WT TP53 solution (**fig. 3m, n**) at a 2% v/v ratio and observed a remarkable seed-dependent growth of WT TP53 fibers (**fig. 3n**). When given sufficient incubation time, a 1:50 mixture of mutant:WT proteins created deposits that were visible to the naked eye (**fig. 3o**). Given the size of these aggregates, these deposits must be constructed almost exclusively from WT proteins, with the mutant proteins serving as the initial seed.

To expand on these observations and ensure that this phenomenon is not caused by artifacts like AFM sample preparation (which involves drying samples on a mica surface), we developed a new hanging drop method to characterize the seeding process in solution (**fig. 4a-c**). First, we prepared a seeding solution of TP53^S149F^ particles with an average size of 0.1 µm as determined by MALS and AFM (**fig. S6**). Then, we set up a 4×6 screening tray with a 1ml reservoir solution that contains the protein buffer and an increasing concentration of NaCl (0.3, 0.5, 0.7, 0.8, 1.0, 1.2M for columns 1-6, respectively). Finally, we added a 10µl WT TP53 solution (60µM) to a siliconized coverslip and placed a 1µl drop of TP53^S149F^ seed particles immediately adjacent to it at different concentrations (0, 1.2, 6, or 12 µM from row A to B, C and D). Over time, the protein drops on the coverslip shrink as a function of the NaCl concentration, gradually increasing the protein concentration. We reasoned that if the mutant seed particles display amyloid potential, this increasing concentration will eventually trigger the conversion of WT proteins to an amyloid state at the drop-drop interface and lead to localized fiber growth (**fig. 4c**). Consistent with this idea, we observed robust growth of TP53 fibers under a light microscope at the WT:mutant interface, but not in the absence of the mutant protein (**fig. 4d-e**). This rod-like material displayed strong birefringence under polarized light, which is highly suggestive of amyloid structures (**fig. 4f**). Taken together, these biophysical experiments strongly support the idea that transcription errors create amyloid proteins that can convert WT proteins to an amyloid state, which initiates the formation of large amyloid fibers and deposits. In addition, they suggest that a limited number of mutant transcripts is sufficient to initiate this process. For example, if 2% of TP53^S149F^ proteins is sufficient to initiate the aggregation of WT TP53, then 2% of TP53 transcripts carrying the TP53^S149F^ error should be sufficient to initiate fiber formation as well. Similar thresholds were previously observed for other amyloid proteins, and it has been speculated that for prions, there may not be a safe dose at all(*44*).

**Fig. 4.**
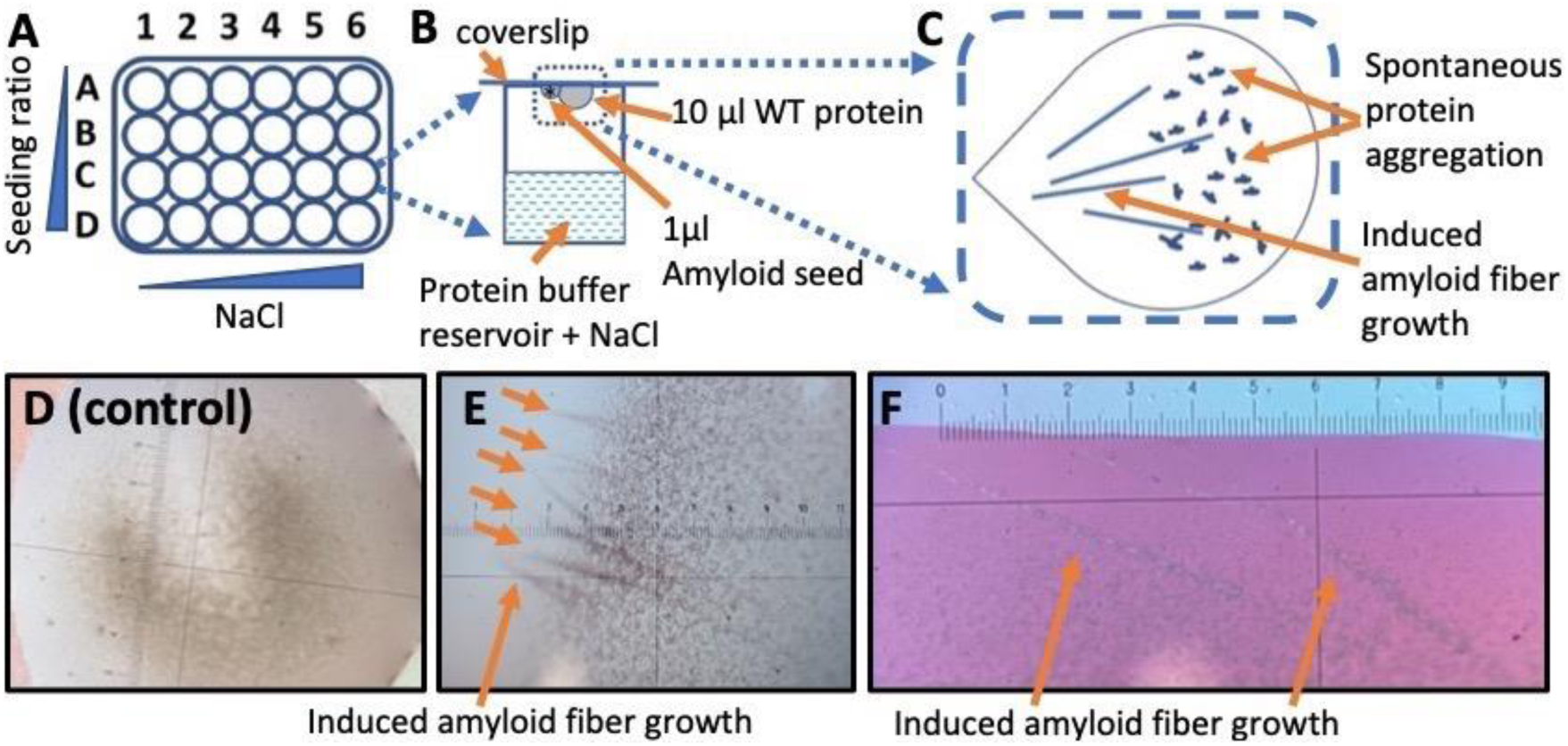
A hanging drop method detects the amyloid and prion-like potential of proteins. **A.** A 4×6 screening tray was set up with a 1ml reservoir that contains protein buffer and an increasing concentration of NaCl. **B.** A 10µl WT TP53 solution (60µM) was then added to a siliconized coverslip and a 1µl drop of TP53^S149F^ seed particles was placed immediately adjacent at decreasing concentrations. **C.** If the mutant seed particles have amyloid potential, this event will trigger conversion of WT proteins at the drop-drop interface and lead to localized fiber growth **D.** If no TP53^S149F^ is provided as seeding material, no fiber-like material forms in the WT TP53 solution. **E.** However, if TP53^S149F^ seeding material is provided, fiber-like material grows out of the WT solution. **F.** These fibers show strong birefringence under polarized light, suggestive of amyloid structures.

With this idea in mind, we decided to test if it is possible for 2% of transcripts to carry the same transcription error. Although this threshold can easily be reached if the number of transcripts generated from a gene is relatively low (<100 per cell), it is less clear if the same is true for highly transcribed genes. Interestingly though, it was previously shown that DNA damage can provoke the same mistake by RNA polymerase II during multiple rounds of transcription(*14*) (**fig. 5a**), so that up to 50% of transcripts can carry the same transcription error(*45–47*). These studies were primarily performed on DNA repair deficient cells though, using a single DNA lesion placed on a plasmid. As a result, it is unclear how well these findings translate to a WT genome carefully wrapped in chromatin that is actively surveyed by DNA repair. Therefore, we designed a new, single cell sequencing approach to examine the impact of DNA damage on transcriptional mutagenesis. First, we treated mouse neuronal stem cells (NSCs) that were derived from the hippocampus for 1 hour with MNNG, a powerful mutagen that randomly generates O_6_-methyl guanine adducts (O_6_-me-G)(*48*). We chose hippocampal stem cells for these experiments because they are directly implicated in amyloid diseases(*49*), and O_6_-me-G adducts because they play an important role in human brain cancers(*50, 51*) and were recently implicated in the pathology of female patients with Alzheimer’s disease(*52*). In addition, we performed these experiments on non-dividing NSCs (**fig. S7**), so that the O_6_-me-G lesions we induced would not be fixed into mutations during DNA replication (a common mechanism to prevent mutations from confounding transcription error measurements(*14, 45–47, 53*)). After MNNG treatment, we provided the cells with fresh medium, and let them recover for increasing periods of time. We then sequenced the transcriptome of single cells at different timepoints (**fig. 5b**) to identify transcription errors that occurred in at least 10% of transcripts from a gene, with a minimum of 40 unique transcripts sequenced. These parameters also prevent direct damage to RNA molecules from affecting our measurements, because it is unlikely that this damage will affect the same nucleotide on multiple RNA molecules. We found that MNNG treatment resulted in a >40-fold increase in transcripts with identical errors after 16 hours of recovery time (**fig. 5c**). The vast majority of these events (which we labeled pseudo-alleles for their ability to generate WT and mutant transcripts) were C→U errors, the most common error induced by O_6_-me-G lesions. Notably, no increase was detected in G→A errors, which would have occurred if O_6_-me-G lesions had been fixed into mutations, demonstrating that our experiment was not confounded by conventional mutagenesis. Consistent with this idea, we found that G→A errors did arise in dividing cells (**fig. S7**). In most cases, pseudo-alleles gave rise to 10-20% of mutant transcripts (**fig. 5d**), greatly exceeding the 2% threshold required for amyloid formation *in vitro*. Consistent with the idea that the error generated by these pseudo-alleles cause protein misfolding and aggregation, we found that treated cells displayed a substantial increase in markers for misfolded proteins and proteotoxic stress, particularly at the timepoint the errors reached their peak (**fig. 5e**). Accordingly, human cells that display error prone transcription(*54*) display increased protein aggregation as well (**fig. S8**). We further note that the number of pseudo-alleles rose over time as more and more genes were transcribed, and were still present 16 hours after exposure, indicating that transcriptional mutagenesis is not only abundant after exposure, but also long-lasting, even in cells capable of DNA repair. Interestingly though, loss of DNA repair is increasing implicated in amyloid diseases(*55*). For example, it was recently reported that the main DNA repair protein for O_6_-me-G lesions in human cells (MGMT(*51*)) is hypermethylated in female patients with AD(*52*), suggesting that in these patients, pseudo-alleles could be present for an extended period of time. To test this hypothesis, we first confirmed that female AD patients indeed display reduced MGMT expression by Western blots (**fig. 5f, S9**). Consistent with a previous study, males did not display this trend (**fig. 5g, S9**). To mimic the impact of reduced MGMT expression on human cells, we deleted the yeast homologue of MGMT (MGT1) in the budding yeast *S. cerevisiae*, arrested them in G1 with α-mating factor (**fig. S10**) and then repeated our experiment with MNNG. Similar to human cells, we found that WT yeast cells displayed an increase in pseudo-alleles immediately after exposure, which declined after DNA repair was able to remove these lesions from the genome (**fig. 5h**). However, in the absence of the MGMT homologue MGT1, the pseudo-alleles remained on the genome, causing transcriptional mutagenesis for an extended period of time. These observations confirm our recent findings, which show that MGT1 removes 90% of DNA lesions within a 6-hour timespan (Vermulst, BioRxiv 2023). Similar to neural stem cells, treated yeast cells displayed increased expression of autophagy genes, molecular chaperones and components of the ubiquitin-proteasome system, indicating that they are under proteotoxic stress (**fig. 5i-l, S11**). Consistent with the idea that these markers are upregulated due to transcript errors, we previously found that yeast cells that display error prone transcription also show increased markers of proteotoxic stress(*13*). In contrast, markers associated with translation (which is inhibited in times of proteotoxic stress) were downregulated (**fig. S12**). MGT1Δ cells showed a prolonged response of these markers, consistent with the prolonged presence of pseudo-alleles on their genome (**5i-k, S12**).

**Figure 5.**
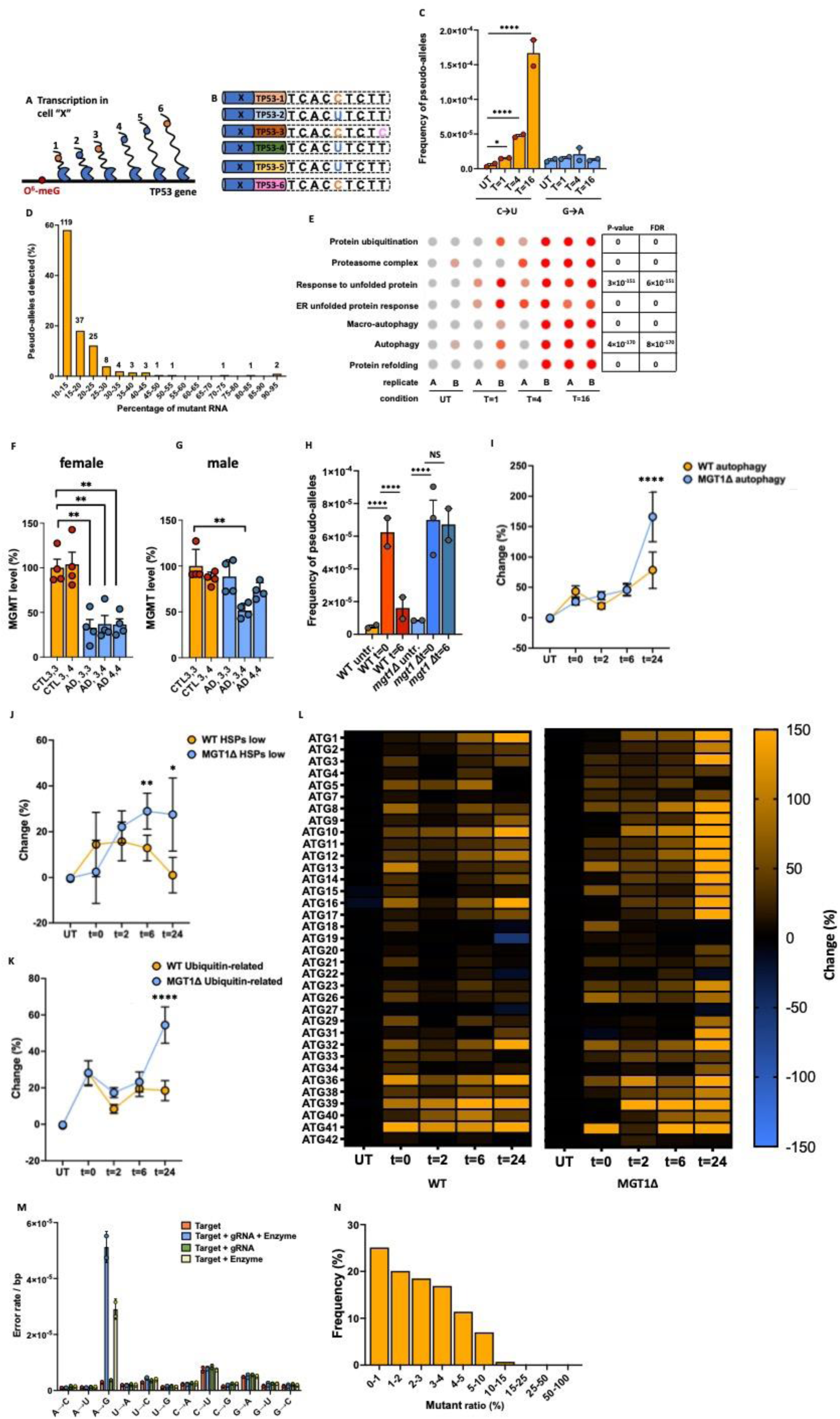
DNA damage and off-target RNA editing affect the fidelity of transcription. **A.** If DNA damage results in multiple rounds of error prone transcription, then multiple transcripts in a single cell should carry identical errors. **B.** When these transcripts are captured and tagged with UMIs, they can be grouped together, and their sequences can be compared to each other to search for identical errors that occur in multiple transcripts. In contrast, sequencing errors or RNA damage will only be present in one transcript. Blue bar: cell-specific barcode. Multicolored bar: transcript UMI. Blue base: WT. Orange base: transcription error. Pink base: Sequencing error/RNA damage **C.** C →U Pseudo-alleles emerge after MNNG treatment created of mouse neuronal stem cells, while G→A errors (which would indicate conventional mutagenesis is occurring as well) do not * = P<0.05, **** = P< 0.0001 according to a Chi-squared test with Yates’ continuity correction. **D.** Ratio of WT:mutant mRNAs identified. Only alleles with more than 10% mutant mRNAs are depicted. **E.** Dot plots of single cell gene expression profiles grouped by GO-terms indicate markers of proteotoxic stress are elevated in treated cells, particularly at 16hours, when the transcript error rate is the highest. Significance was ascertained by ANOVA test. FDR= False Discovery Rate. **F.** MGMT levels are decreased in all females with AD ** = P<0.01, unpaired t-test with Welch’s correction. **G.** MGMT levels are not decreased in males with AD, except for those with a APOE3/APOE4 genotype. **H.** Loss of MGT1, the yeast homologue of MGMT allows O^6^-me-G lesions to remain on the genome, resulting in greatly increased numbers of pseudo-alleles over time. **** = P< 0.0001 according to a Chi-squared test with Yates’ continuity correction. Consistent with the idea that these errors result in misfolded proteins, these cells displayed markers of proteotoxic stress, including upregulated autophagy genes (**I**), heat shock proteins (**J**) and proteins implicated in the ubiquitin-proteasome system (**K**). Depicted in in figure **I** and **K** is the average percentage change for all autophagy and ubiquitination-related genes identified by bulk RNA-seq. The genes depicted in **J** have been separated from several heat shock proteins that displayed unusually large increases in transcript levels (**fig. S9**). * = P<0.05, ** = P<0.01, **** = P< 0.0001 according to a paired t-test. **L.** Heat map of autophagy genes detected in WT and mutant cells. **M.** Error spectrum of human cells after transformation with plasmid that carries an editing target, the gRNA required to edit the target, and the editing enzyme. If the editing enzyme is present, large numbers of A to I editing events (A to G errors) were observed. **N.** Percentage of editing events that generate mRNAs with various mutant:WT ratios. Error bars indicate standard error of the mean.

In addition to transcription errors, it has been proposed that other molecular mistakes could result in amyloid and prion-like proteins as well, including off-target RNA editing(*15*). One of the best-known examples of RNA editing in the animal kingdom is seen in cephalopods(*56*) where ADAR1 edits adenine to inosine (A to I) in a sequence-specific manner, an event that can be monitored by circ-seq as A →G errors(*19*). If off-target editing indeed results in mutant RNAs, then squid and octopi tissues that express high levels of ADAR1 should display high levels of off-target editing, while tissues that do not express ADAR1 should not. Consistent with this idea, we found that the optic lobe and stellate ganglia of cephalopods (which express high levels of ADAR1) displayed large amounts of off-target editing, while the gills (which express low levels of ADAR1) did not (**fig. S13**). These observations suggest that off-target editing is indeed one additional mechanism by which erroneous transcripts can be created. This possibility is particularly important from a medical perspective, as RNA editing tools are increasingly thought of as a new tool to treat symptoms of disease(*57*). To test whether RNA editing tools designed in the lab can result in off-target editing as well, we expressed an RNA editor specifically designed to edit the ATP1a3 transcript in human cells and monitored off-target editing. Similar to our observations in cephalopods, we found that these editors display a substantial amount of off-target editing, whether a guide RNA is present or not (**fig. 5m**). These events resulted in large numbers of rare (<2%) and common (>2%) mutant RNAs (**fig. 5n**), suggesting that these editors have the potential to induce protein aggregation in human cells. These observations suggest that all RNA editors designed for clinical purposes should go through rigorous testing prior to use, and that circ-seq is a highly sensitive tool to detect these potentially deleterious side effects.

## DISCUSSION

To identify the molecular mechanisms that underpin human aging and understand how these mechanisms drive age-related pathology, it will be essential to determine how amyloid and prion-like proteins are generated. Here, we demonstrate that transcript errors represent one of these mechanisms. Although transcript errors are transient events, amyloid and prion-like proteins are characterized by their ability to “replicate themselves” by converting WT proteins to an amyloid state. Thus, even a transient event like a transcription error could generate enough mutant proteins to trigger this reaction.

One of the most intriguing observations from our experiments is that transcript errors generate mutant proteins that are already known to cause familial cases of amyloid disease. Thus, our experiments raise the remarkable possibility that both genetic and non-genetic cases of amyloid diseases could be caused by identical mutant proteins, only the mechanism by which they are generated is different (**fig. 6**). Interestingly, it was previously found that aggregates of tau have identical structures in both familial and non-familial cases of AD (*58*), suggesting that they were initiated by identical mutant proteins.

**Figure 6.**
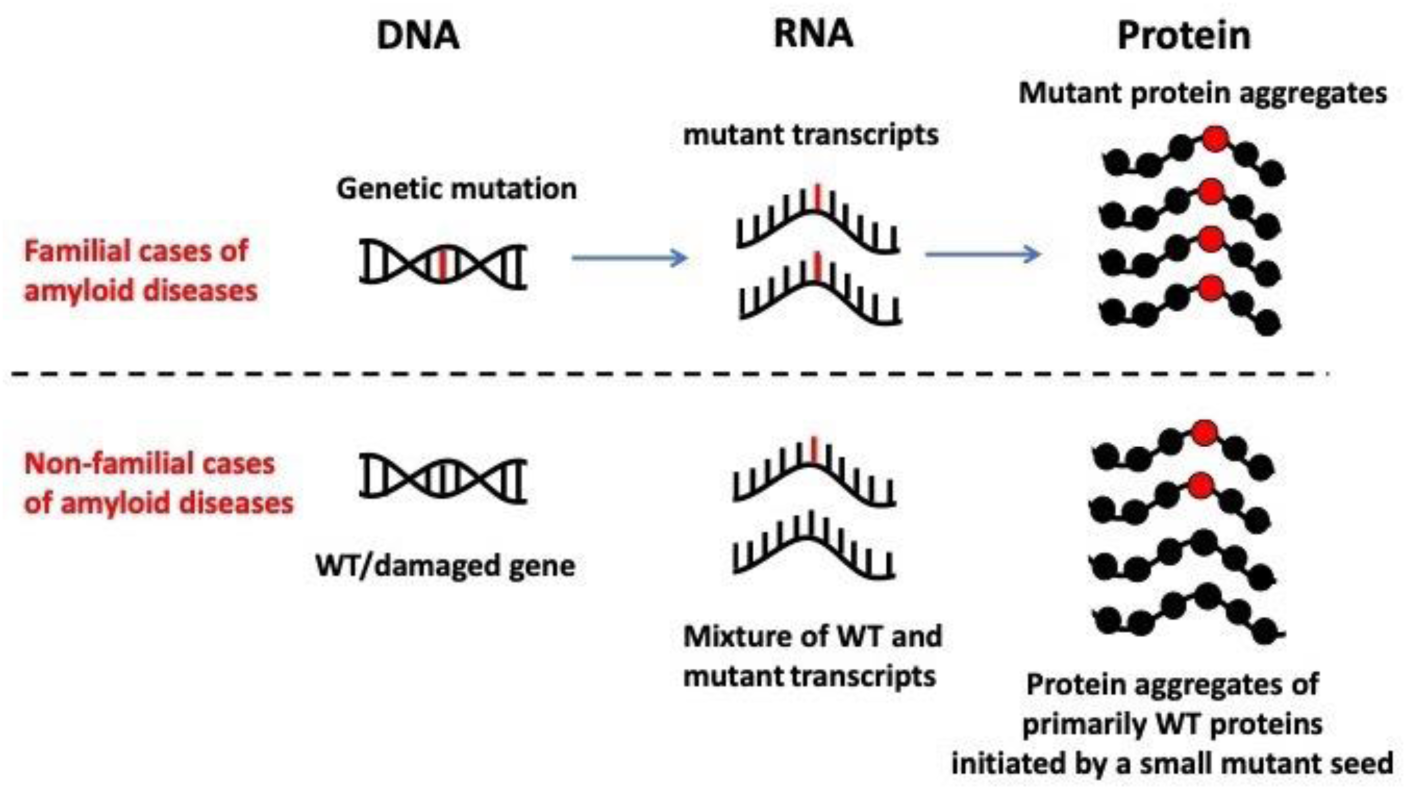
Model for the contribution of transcription errors to non-familial cases of disease. In familial cases of amyloid diseases, genetic mutations generate mutant protein with greatly increased amyloid potential. In non-familial cases, identical mutant proteins, as well as unique ones, may be generated by transcription errors.

Transcription errors also generate amyloid proteins that have not been characterized yet. The mutant version of TP53^S149F^ we examined here is one example of this phenomenon. For example, we recently identified a number of additional novel mutant proteins that seem to have similar features compared to TP53^S149F^ (data not shown). We conclude therefore, that in addition to the mutant proteins already known to play a role in amyloid diseases, transcription errors also generate amyloid proteins that have thus far escaped detection. However, these proteins are likely to have the same potential to affect cellular proteostasis and induce protein aggregation.

Our quantitative experiments further show that 1-2% of TP53^S149F^ is sufficient to initiate aggregation of WT TP53. Remarkably, this threshold is easily breached with the help of DNA damage, a form of cellular stress that has long been associated with protein misfolding diseases. For example, farmers that are exposed to the DNA damaging pesticides rotenone and paraquat have an increased risk for developing Alzheimer’s disease and Parkinson’s disease(*59, 60*), while the DNA damaging agent methylazoxymethanol (MAM) is suspected of being the pathogenic agent responsible for ALS/Guam-Parkinsonism-Dementia Complex, a disease that is characterized by protein aggregation and dementia-like symptoms(*61–63*). We recently discovered that MAM induces pseudo-alleles in mouse neural stem cells as well (Verheijen, bioRxiv 2023). Finally, DNA damage is ubiquitous feature of aging cells the primary risk factor for protein misfolding diseases. Our data, and data by others(*45–47*), provides a compelling rationale for these observations by demonstrating that DNA damage creates long-lasting pseudo-allele across the genome that give rise to a mixture of WT and mutant transcripts. As a result, 10-30% of the transcripts can carry the same transcription error, a ratio that greatly exceeds the amount required to initiate aggregation. Although our experiments focused on pseudo-alleles that were created by O^6^-me-G, it should be noted that other forms of DNA damage can generate pseudo-alleles as well(*45–47*), including oxidative DNA damage(*14*). Thus, other aspects of human aging that are known to produce oxidative damage (including mitochondrial dysfunction and inflammation) could trigger protein aggregation through a similar mechanism. A similar rationale applies to environmental factors such as pollution and lifestyle choices such as smoking.

Remarkably, we found that after one treatment of MNNG, approximately 1 out of every 6,000 guanine bases was converted into a pseudo-allele, which is expected to affect ~1 out 10 genes. Thus, one exposure to a DNA damaging agent could result in thousands of pseudo-alleles emerging across the genome, demonstrating the potential of DNA damage to generate large amounts of identical mutant proteins without the need to induce mutations: the damage itself is sufficient.

Consistent with a role for DNA damage in protein misfolding diseases, it is now increasingly recognized that loss of DNA repair can exacerbate amyloid diseases as well. For example, DNA repair is thought to play an important role in ALS(*55*), and it was recently shown that the DNA repair gene MGMT is hypermethylated in female patients with AD(*52*). When we mimicked this phenomenon in yeast, we found that reduced DNA repair allows pseudo-alleles to persist on the genome for extended periods of time, creating vast amounts of mutant proteins and a prolonged presence of markers associated with loss of proteostasis. It has long been known that females are at a greater risk for AD compared to males, and our data suggests that reduced MGMT expression, followed by extended transcriptional mutagenesis, could help explain this sex-specific difference(*64*).

Besides DNA damage, the fidelity of transcription can also be altered by other variables. For example, we previously found that the error rate of transcription increases with age in yeast(*13*) and flies, and is affected by epigenetic markers, cell type and genetic context(*12, 19*). Each of these variables could alter the rate with which amyloid proteins are generated. One important example of this idea was observed in patients with non-familial cases of Alzheimer’s disease. In these patients, transcription errors occur on dinucleotide repeats in the APP and the UBB gene, two key proteins associated with Alzheimer’s disease. These errors generated shortened peptides that were later found to be present in the amyloid plaques that characterize the disease, suggesting that they play a role in pathogenesis(*65, 66*). We recently demonstrated that repetitive DNA sequences can increase the error rate of transcription up to a 100-fold in human cells, directly supporting this observation. Moreover, we used a novel mouse model to demonstrate that neurons in the hippocampus are especially prone to making these transcription errors, providing further evidence for a link between transcriptional mutagenesis and Alzheimer’s disease(*19*).

Finally, it is important to note that it may not be necessary for transcription errors to create highly specific amyloid proteins to promote amyloid diseases. Surprisingly, we previously found that random transcription errors can affect protein aggregation as well. Because the primary impact of mistakes in protein coding sequences is protein misfolding(*67*), random errors tend to create a cache of misfolded proteins that affect the entire proteome. Although most of these misfolded proteins are benign, and do not affect the aggregation of disease related proteins directly, they do need to be degraded by the same protein quality control machinery. As a result, random errors can create enough misfolded proteins to overwhelm the protein quality control machinery, which then allows pathological amyloid proteins to evade degradation and seed aggregates(*13*). Thus, transcription errors may not only generate highly specific amyloid and prion-like proteins, as we demonstrate here, they may also generate the very conditions that allow these proteins to evade the protein quality control machinery and initiate aggregation.

## FIGURES

**Figure S1.**
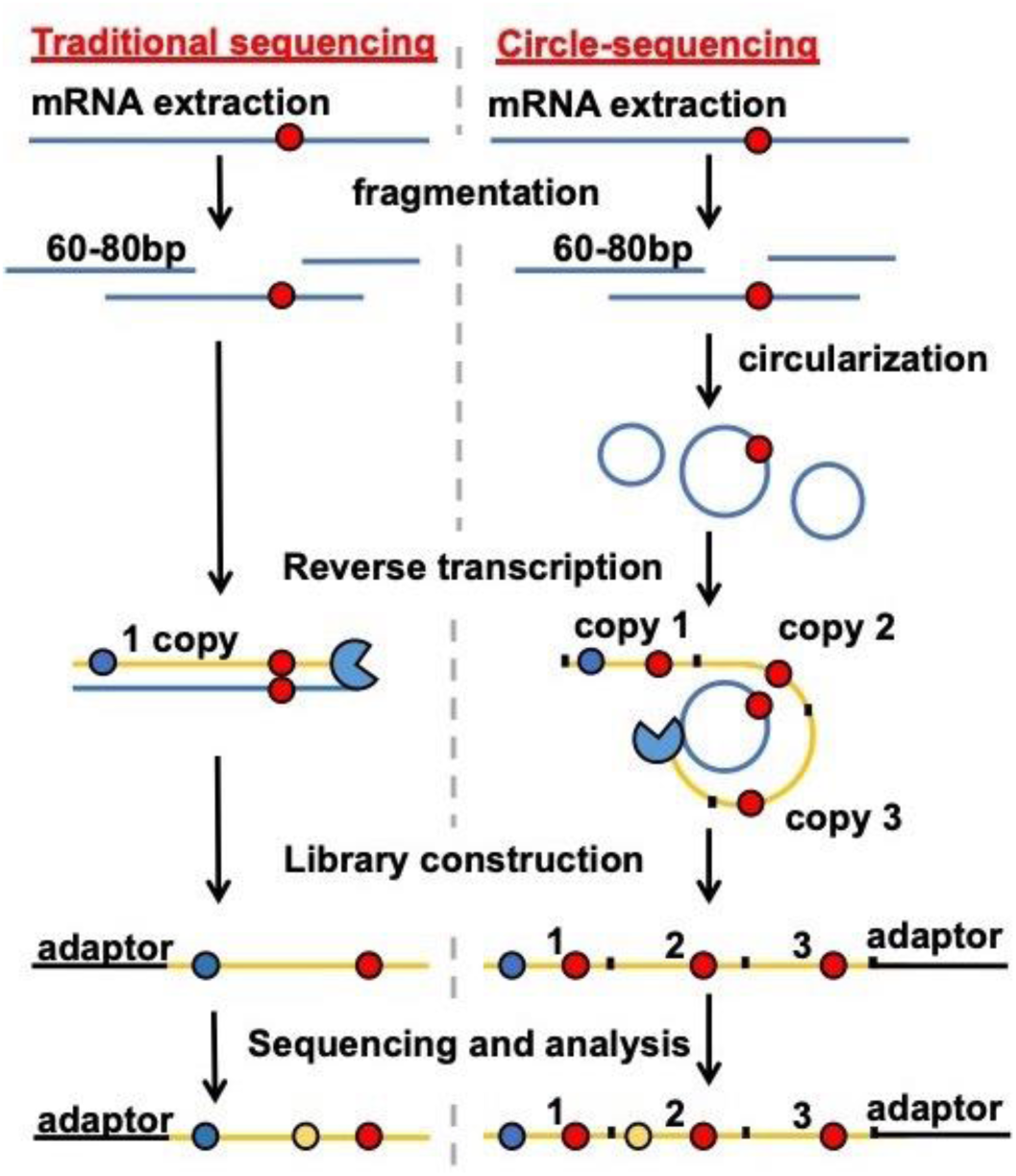
Concept of the circle-sequencing assay. Data from traditional cDNA libraries are littered with RT (blue circles) and sequencing errors (orange circles) that are indistinguishable from true transcription errors (red circles). To remove these artifacts from our data, we circularize RNA (blue lines) prior to RT. These circularized molecules are then reverse transcribed in a rolling circle fashion to generate linear cDNA molecules made from tandem repeats of the template. If an error was present in the template, that error will be present in all copies of the cDNA molecule (yellow lines), while artifacts are present in only one.

**Figure S2.**
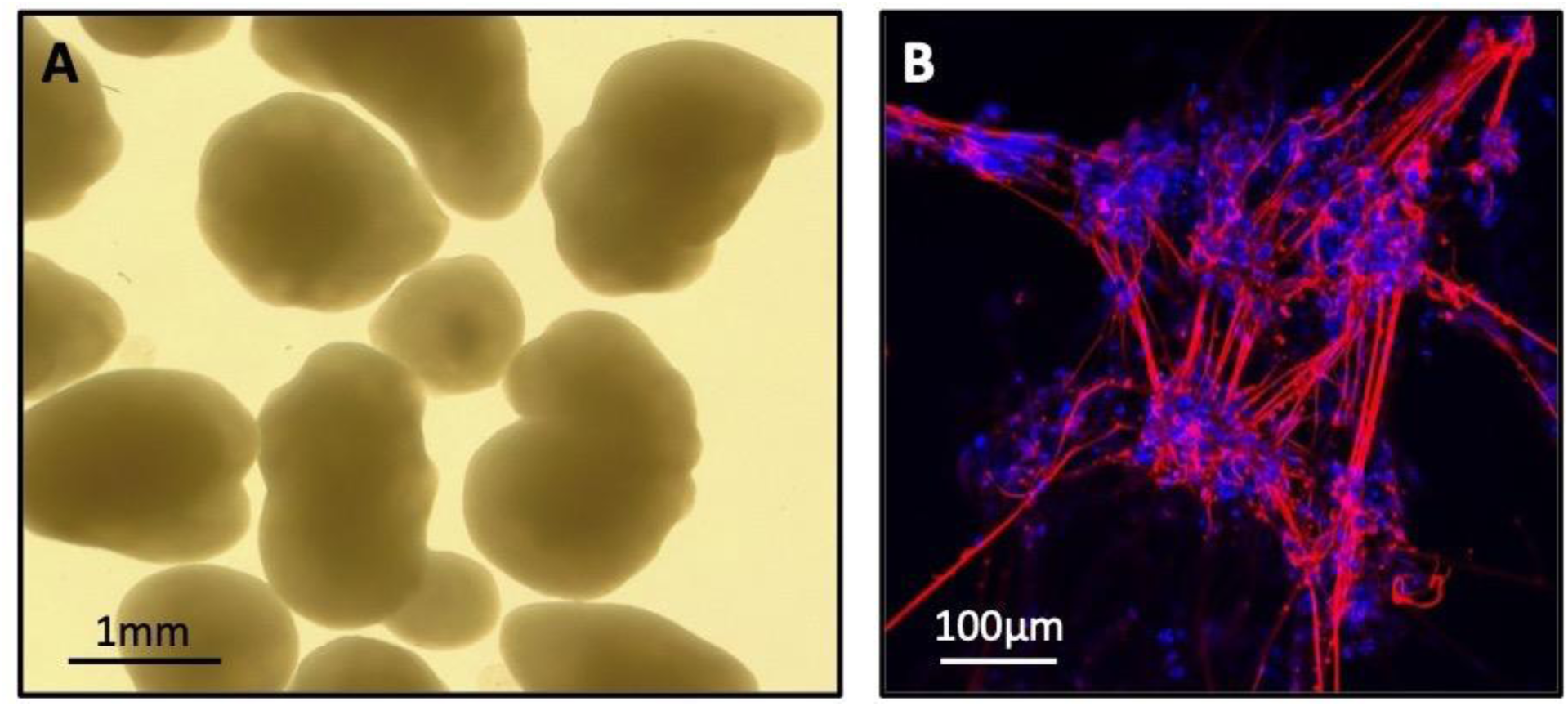
brain organoids and neurons created from H1 ESCs. **A.** Brain organoids, visible by the naked eye, were generated from H1 ESCs. Approximately 60 organoids were generated for each biological replicate and cultured for 3 months to mature them. **B.** H1 ESC-differentiated Ngn2-neurons stained for DAPI and βIII-tubulin, a pan-neuronal cytoskeletal marker showing neuronal grow projections.

**Table S1.**
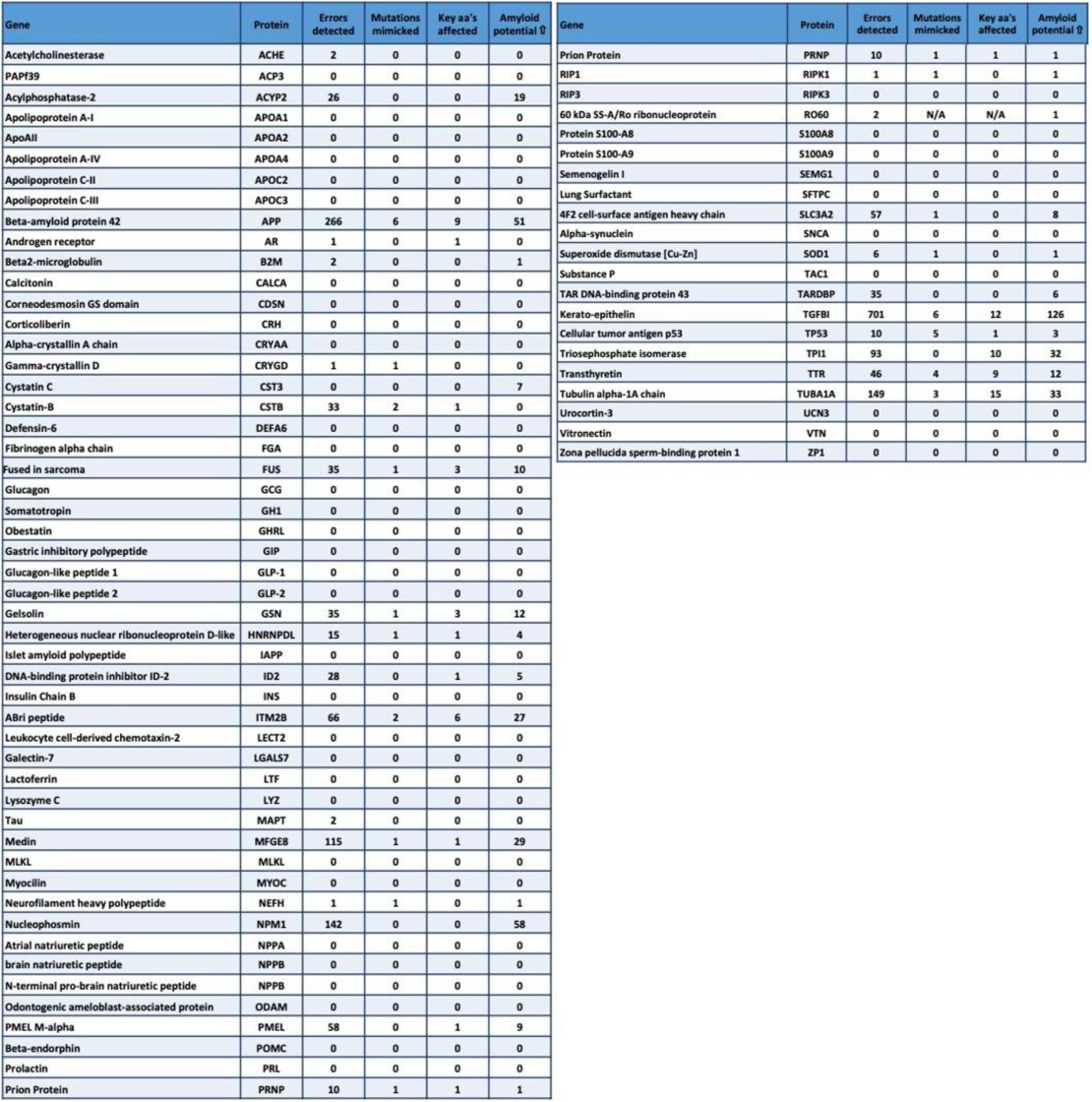
Errors detected in transcripts that code for amyloid proteins known to be implicated in disease, as well as proteins with amyloid cores. **Column 1:** Gene name. **Column 2:** Protein name. **Column 3:** Number of errors detected in transcripts that code for this protein. **Column 4:** Number of errors that generate the same mutant proteins as seen in familial cases of disease. **Column 5:** Number of errors that affect the same amino acid (aa) as affected in disease, but change it to a different residue compared to the clinic. **Column 6:** Number of errors that increase the amyloid potential of these proteins as predicted by bio-informatic analysis (AmyPred-FRL).

**Table S2.**
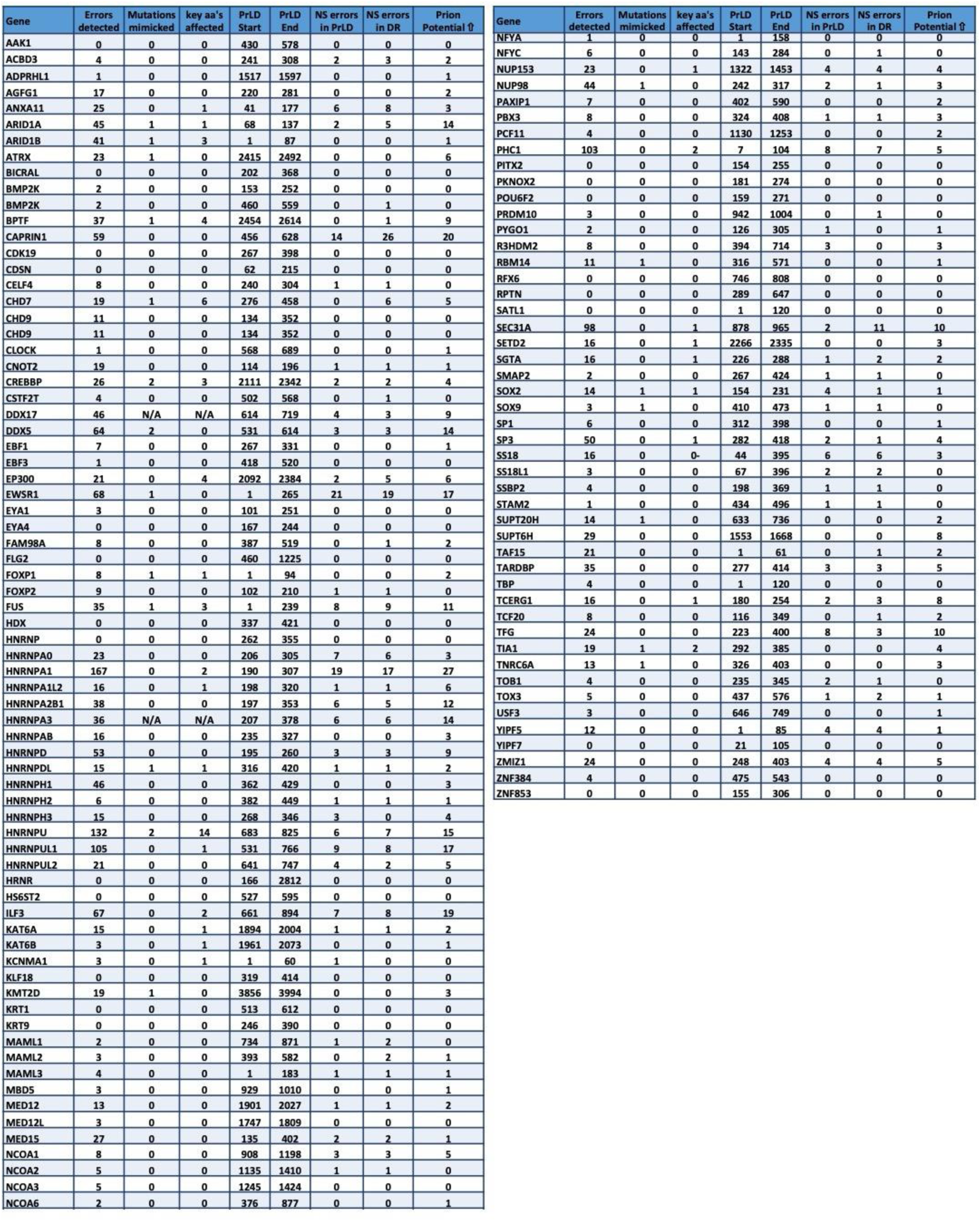
Errors that affect the prion-like domain of proteins. **Column 1:** Gene name. **Column 2:** Number of errors detected in transcripts that derived from this gene. **Column 3:** Number of errors that generate mutant proteins identical to those seen in familial cases of prion-like diseases. **Column 4:** Number of errors that affect an amino acid (aa) known to be involved in disease, but mutate it to a different residue compared to the clinic. **Column 5:** Start of prion like domain (PrlD) in protein. **Column 6:** End of prion like domain in protein. **Column 7:** Number of non-synonymous errors that affect the prion-like domain. **Column 8:** Number of non-synonymous errors that affect the disordered region of the protein. **Column 9:** Number of errors that increase the prion-like potential of the protein as predicted by bio-informatic analysis (PAPA).

**Figure S3.**
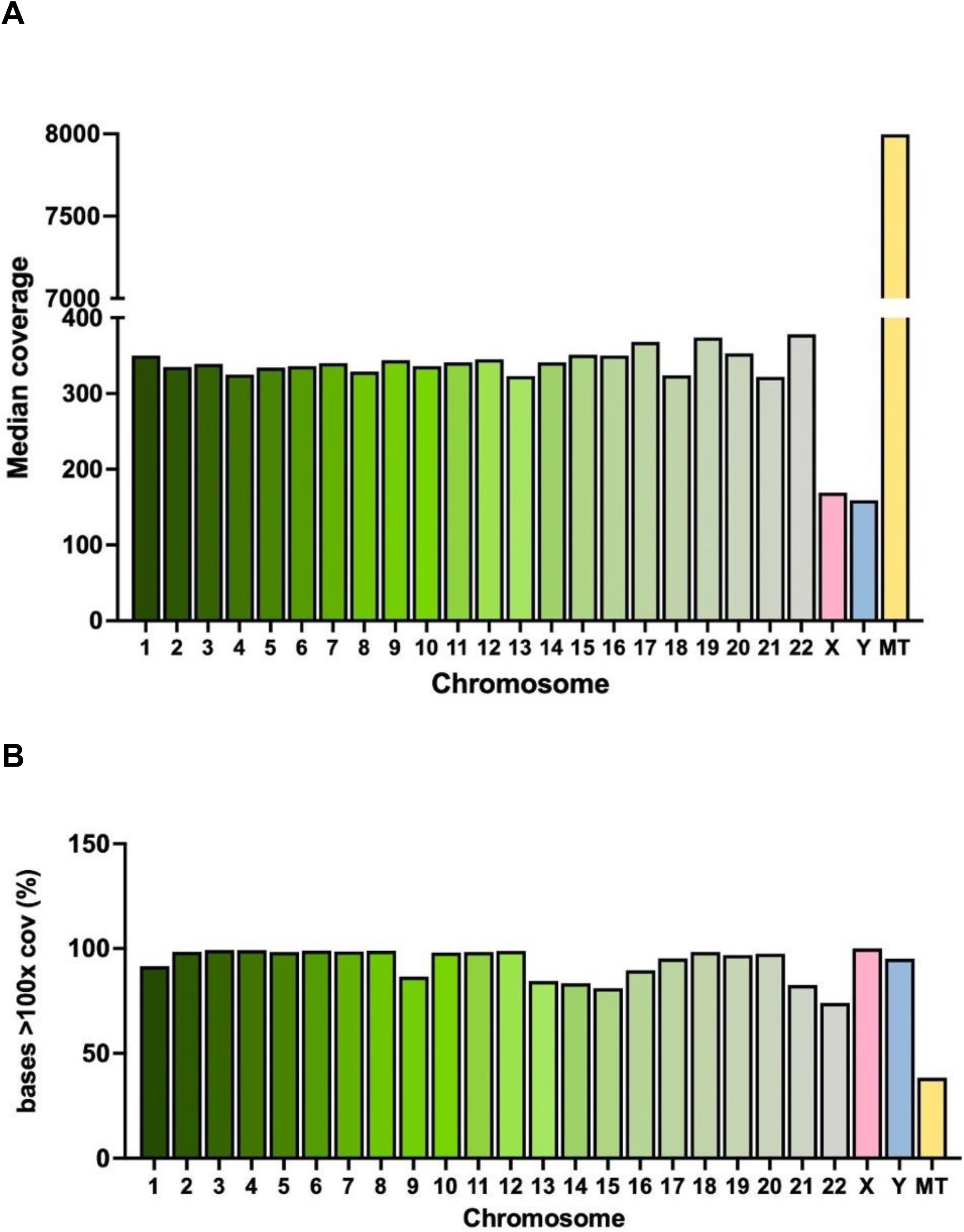
Median coverage of chromosomes in H1 ESCs cells by DNA-seq. **A.** Median coverage across all chromosomes. **B.** Percentage of bases that were covered at least 100x. This data was used to create a reference genome for the H1 ESCs and the brain organoids and neurons generated from these cells.

**Figure S4.**
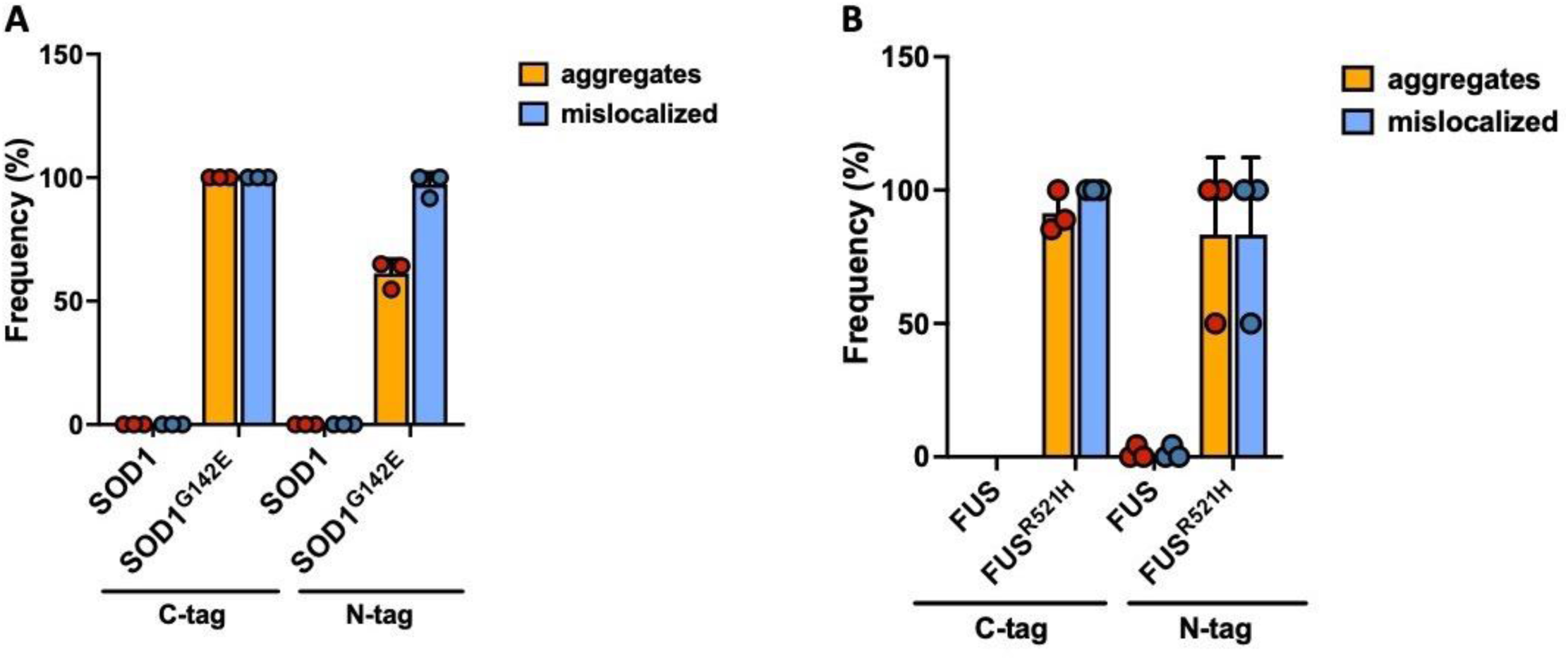
Amyloid behavior of SOD1 and FUS in HEK293 cells. In addition to primary fibroblasts, SOD1 and FUS also aggregate and mislocalize in HEK293 cells. Depicted are the % of cells with aggregates or mislocalized proteins. Error bars indicate standard error of the mean.

**Figure S5.**
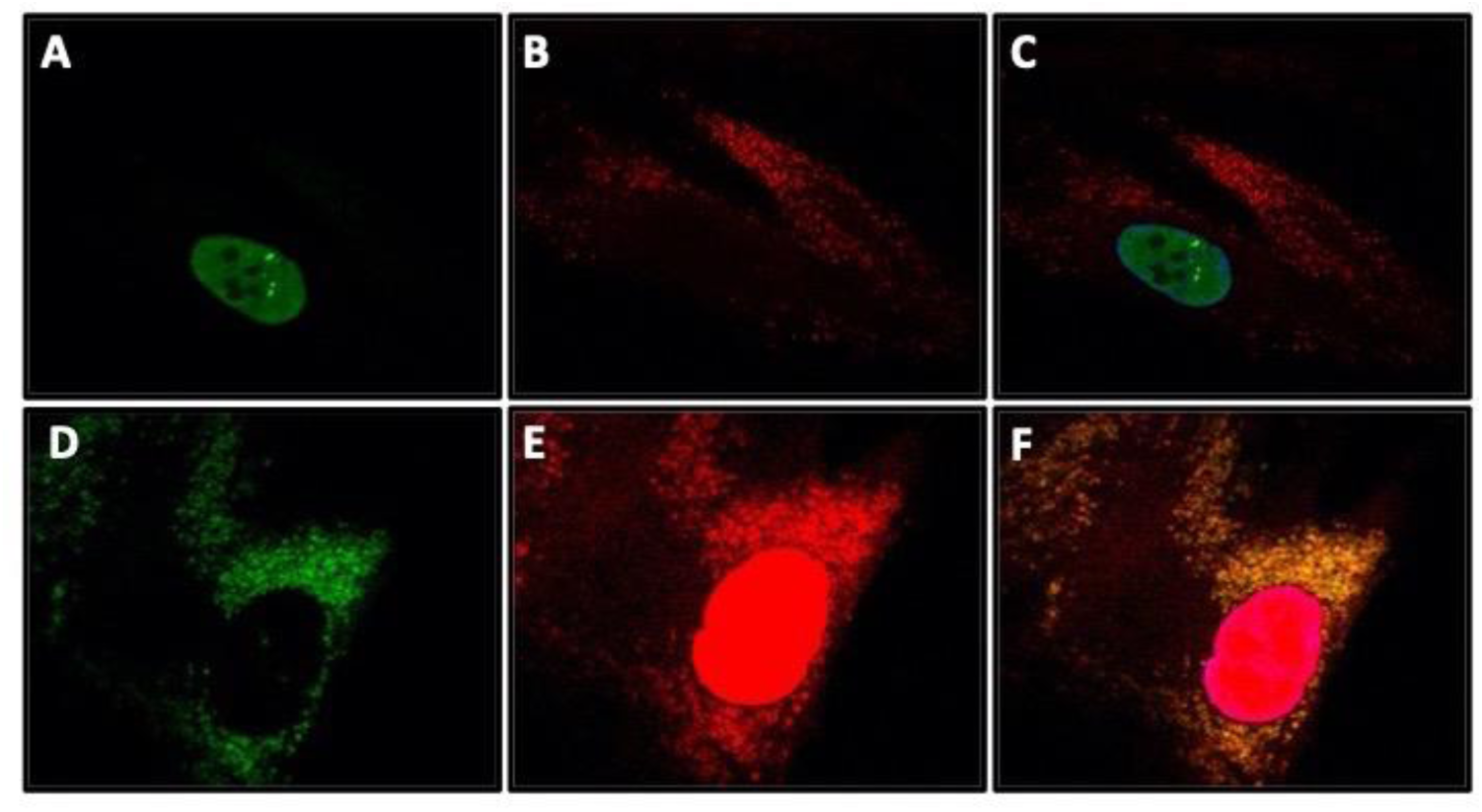
FUS aggregation in primary fibroblasts. **A-C.** WT FUS (green) almost always colocalized with mutant FUS^R521H^ in the cytoplasm (red), although rare exceptions were detected. **D-F.** Mutant FUS^R521H^ is almost always present in the cytoplasm, although rare exceptions were detected where the nucleus was filled with FUS^R521H^. However, in those cases, FUS^R521H^ did not form clear, functional foci similar to WT FUS.

**Figure S6.**
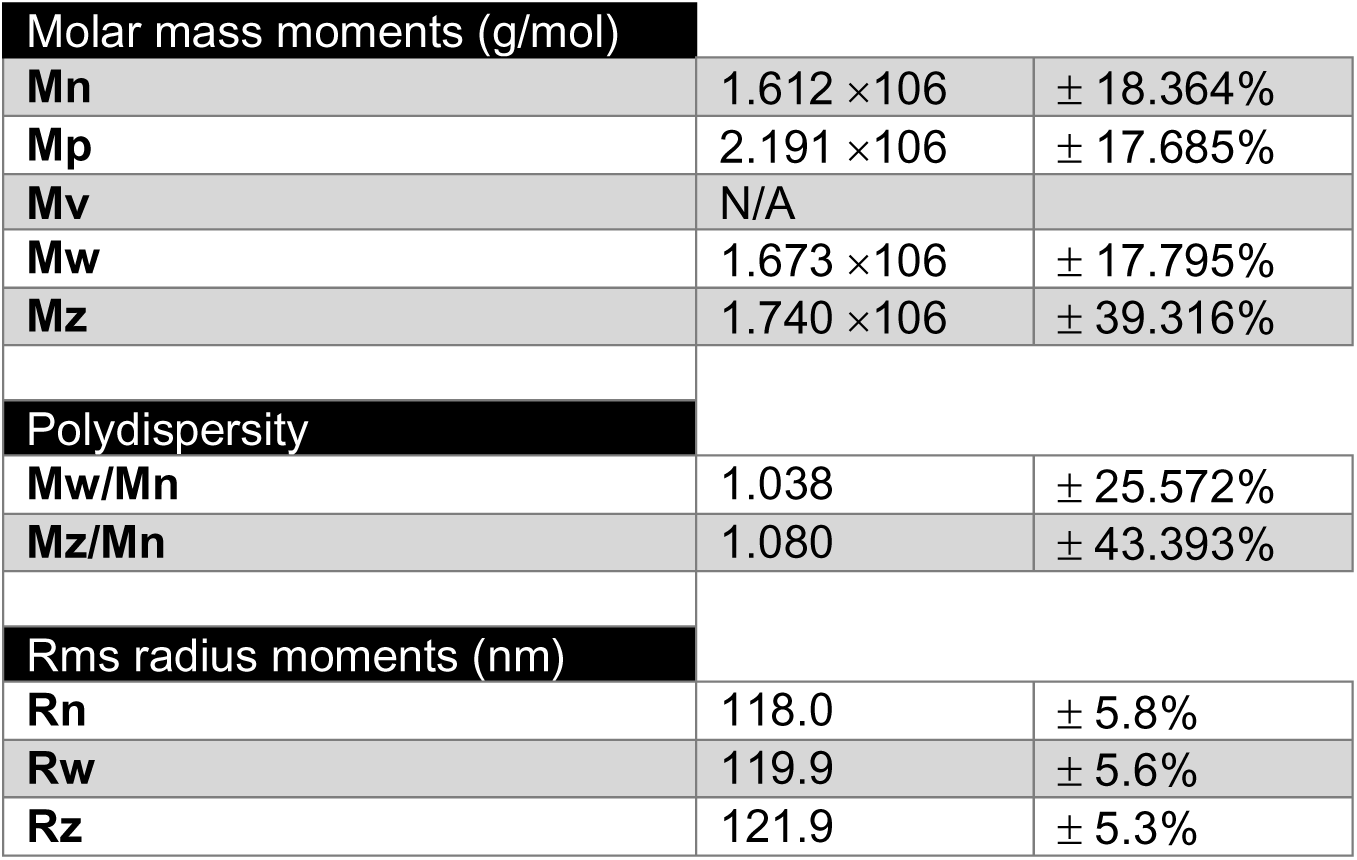
MALS based calculations of TP53^S149F^ particle size.

**Figure S7.**
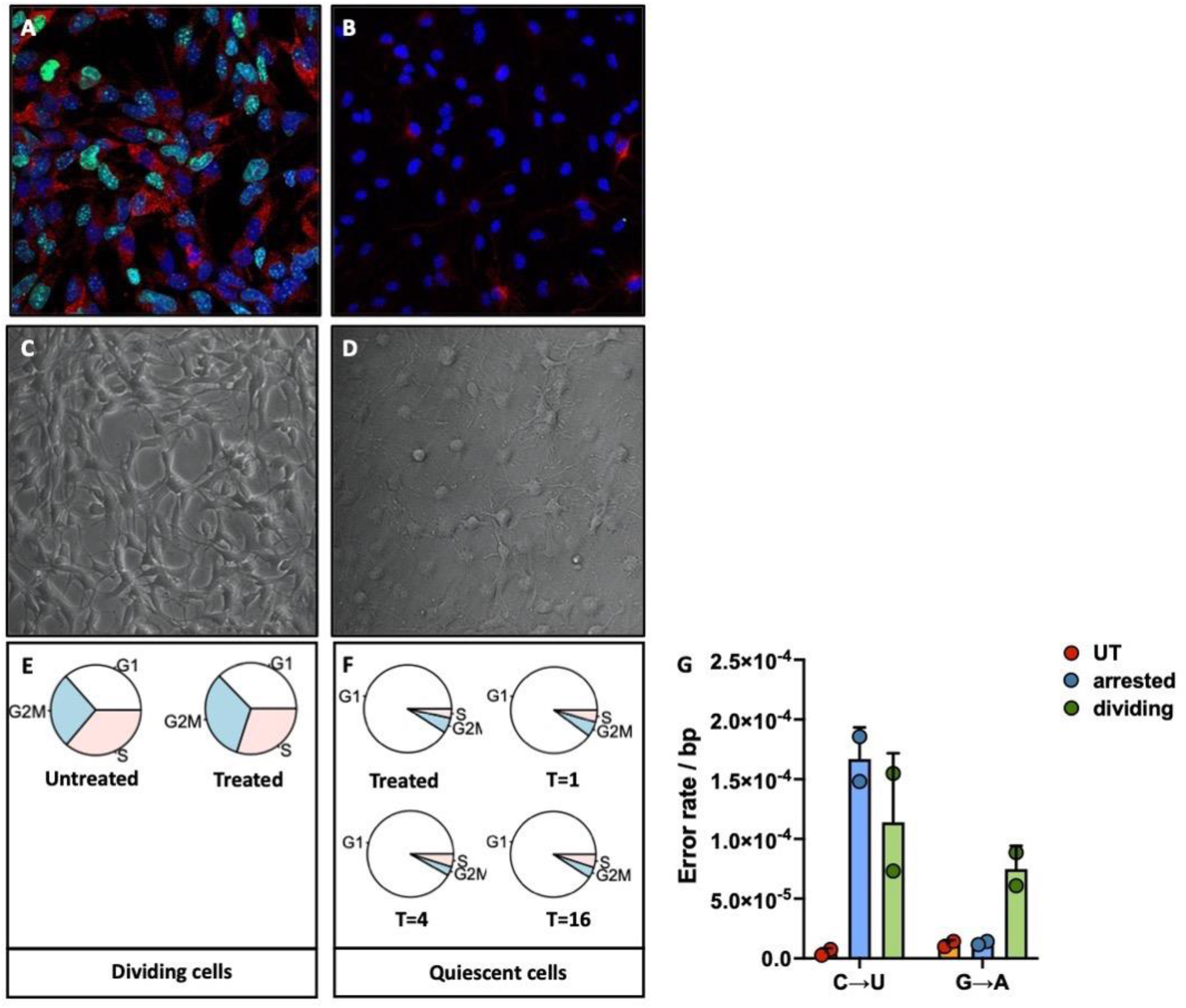
Mouse neural stem cells brought into a quiescent state. Mouse neural stem cells were either cultured in a proliferating or a quiescent state. Proliferation was assessed by Ki67 staining, cell size and shape and transcriptomic analysis. **A.** Proliferating mNSCs show bright Ki67 staining (green), while cells brought into quiescence by addition of bmp4 to the cell culture medium were not (**B**). **C.** Cells in a proliferating state show a distinct difference in size and shape compared to cells brought into a quiescent state (**D**). **E-F.** Single cell transcriptome analysis of dividing and quiescent mouse neural stem cells used in this study. **G.** Quiescent cells treated with MNNG show only an increase in C→U errors, indicative of transcription errors made on O^6^-meG lesions, while dividing cells also displayed an increase in G→A errors indicating that they have undergone conventional genetic mutagenesis as well. Error bars indicate standard error of the mean.

**Figure S8.**
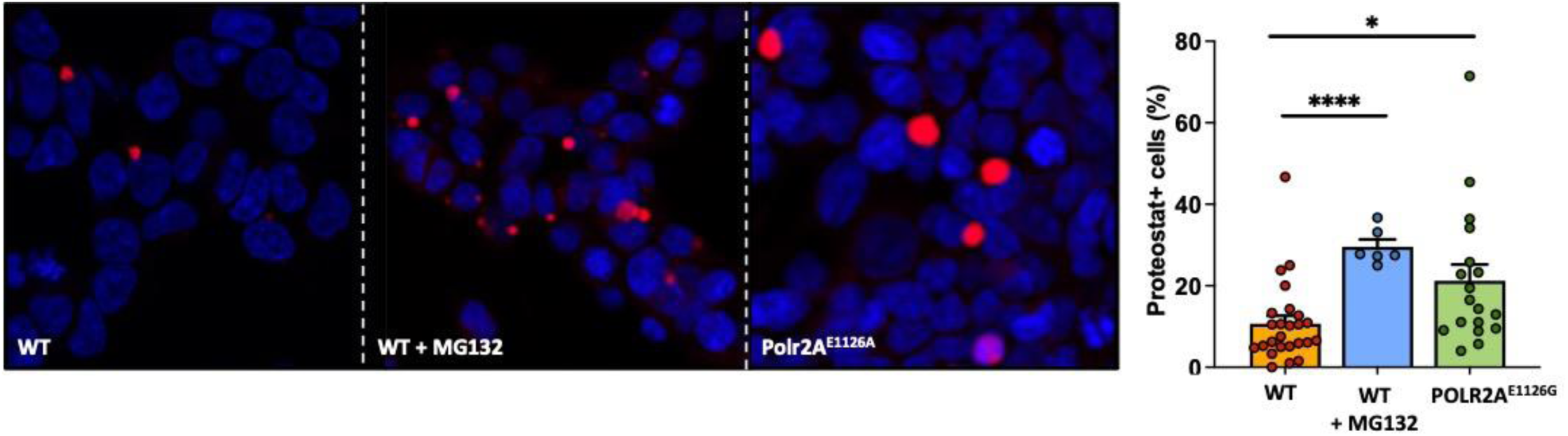
Proteostat staining of WT and Polr2AE1126G cells. **A.** WT cells stained with proteostat, a dye that highlights protein aggregates. **B.** WT cells treated with MG132, which blocks the proteasome, leading to the accumulation of misfolded proteins and protein aggregates. **C.** Polr2AE1126G cells display a 3-fold increase in transcription errors. Consistent with the idea that transcription errors cause protein misfolding and protein aggregation, these cells display increased staining for proteostat.

**Figure S9.**
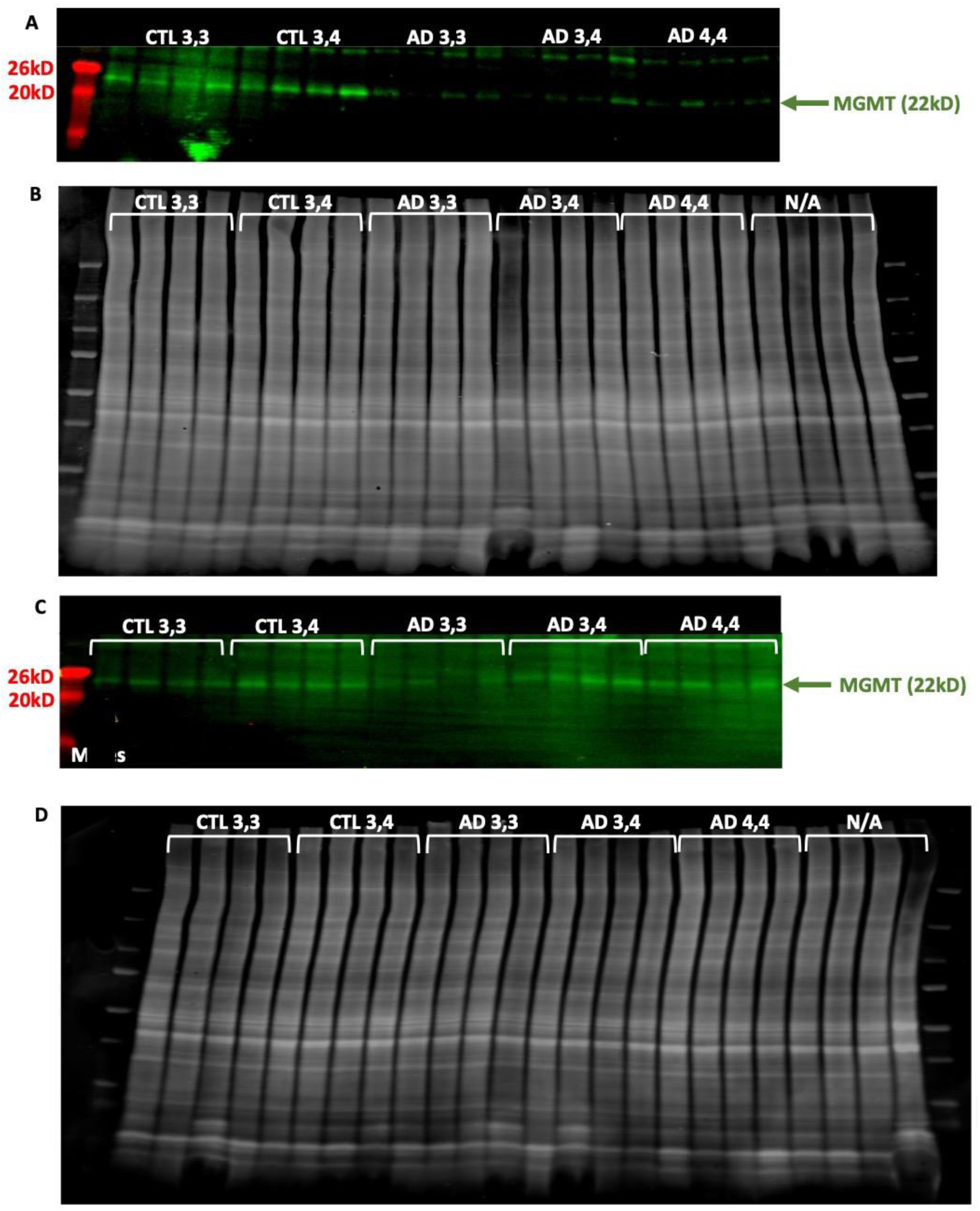
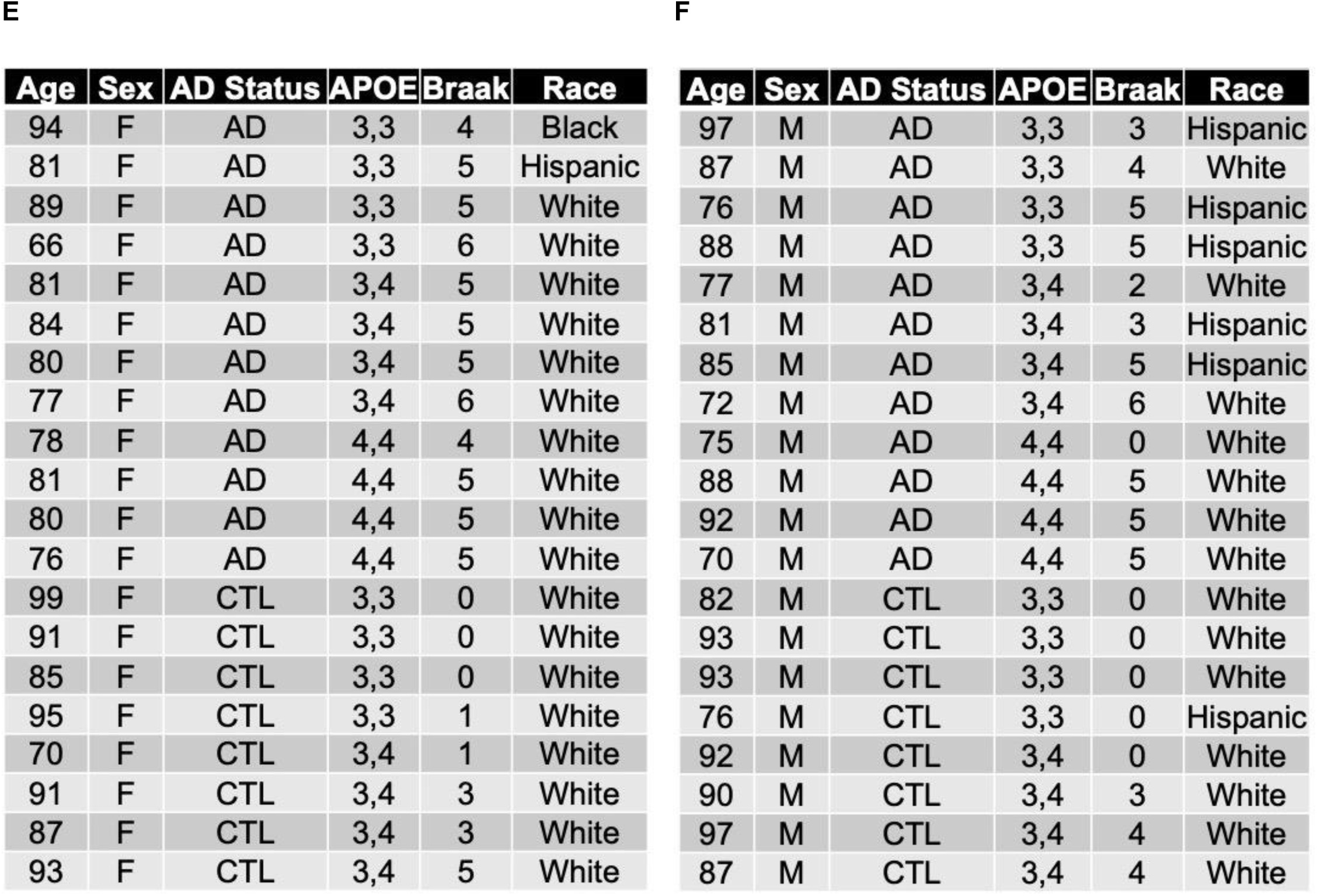
MGMT levels in female and male AD patients and carriers with or without an APOE4 allele. **A.** Western blot (**A, C**) used to quantify the levels of MGMT in females (**fig. 5e**) and males (**5f**). Ponceau staining (**B, D**) was used for normalization. N/A indicates samples not used for this study**. E-F.** APOE status of patients and controls used for **figure 5e** and **S8A-D.**

**Figure S10.**
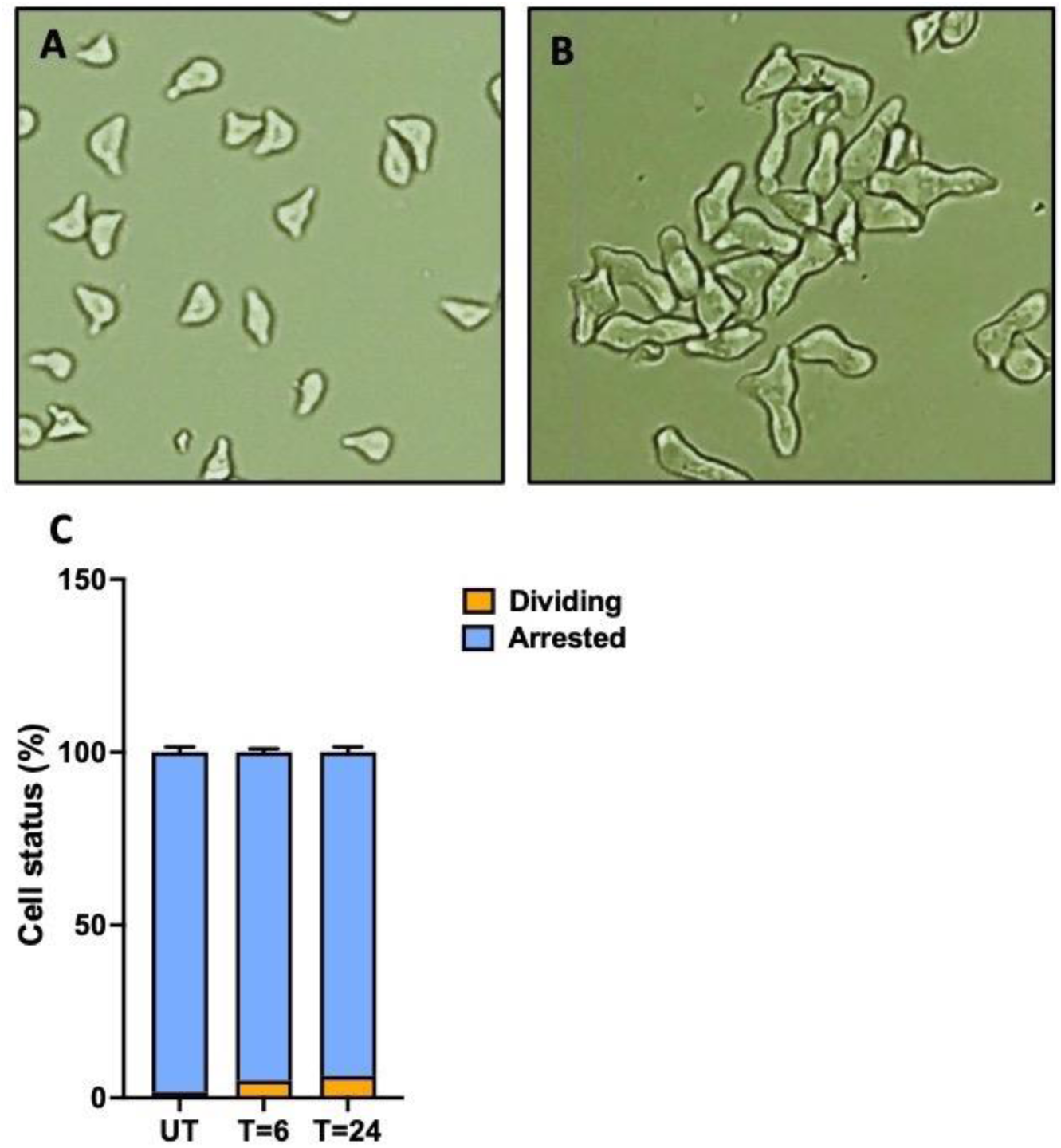
Cell cycle arrest of yeast cells with α-mating factor. Cells were arrested with 50ng/ml α-mating factor for 6 hours, resulting in characteristic pear-shape cell structures (**A**). After 6 hours, another 50ng/ml was added to the cells for overnight incubation leading to further alterations in cell structure (**B**). **C.** Cell cycle arrest was quantified by counting actively budding yeast cells and cells with canonical pear-shape structure induced by α-mating factor. Error bars indicate standard error of the mean.

**Figure S11.**
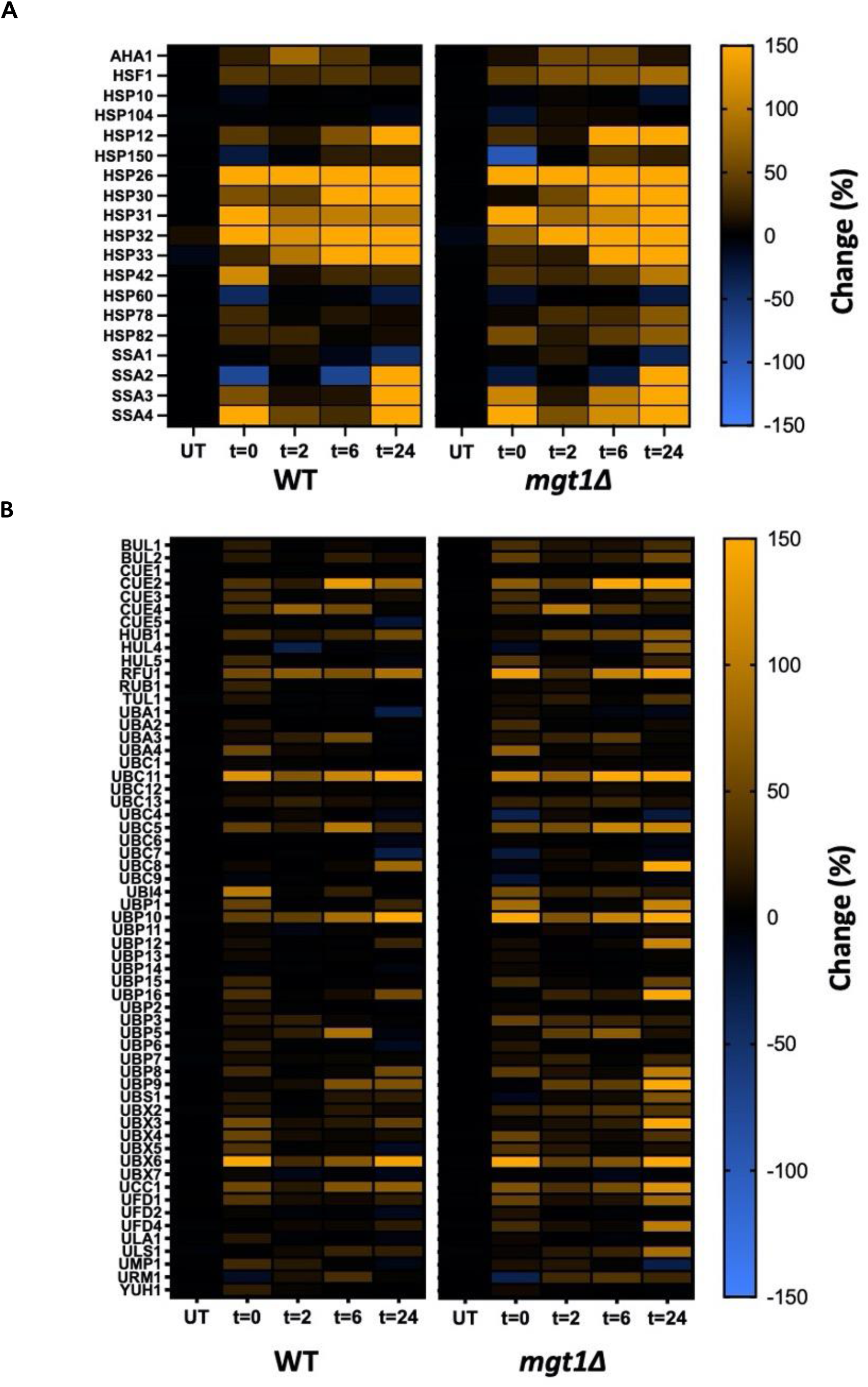
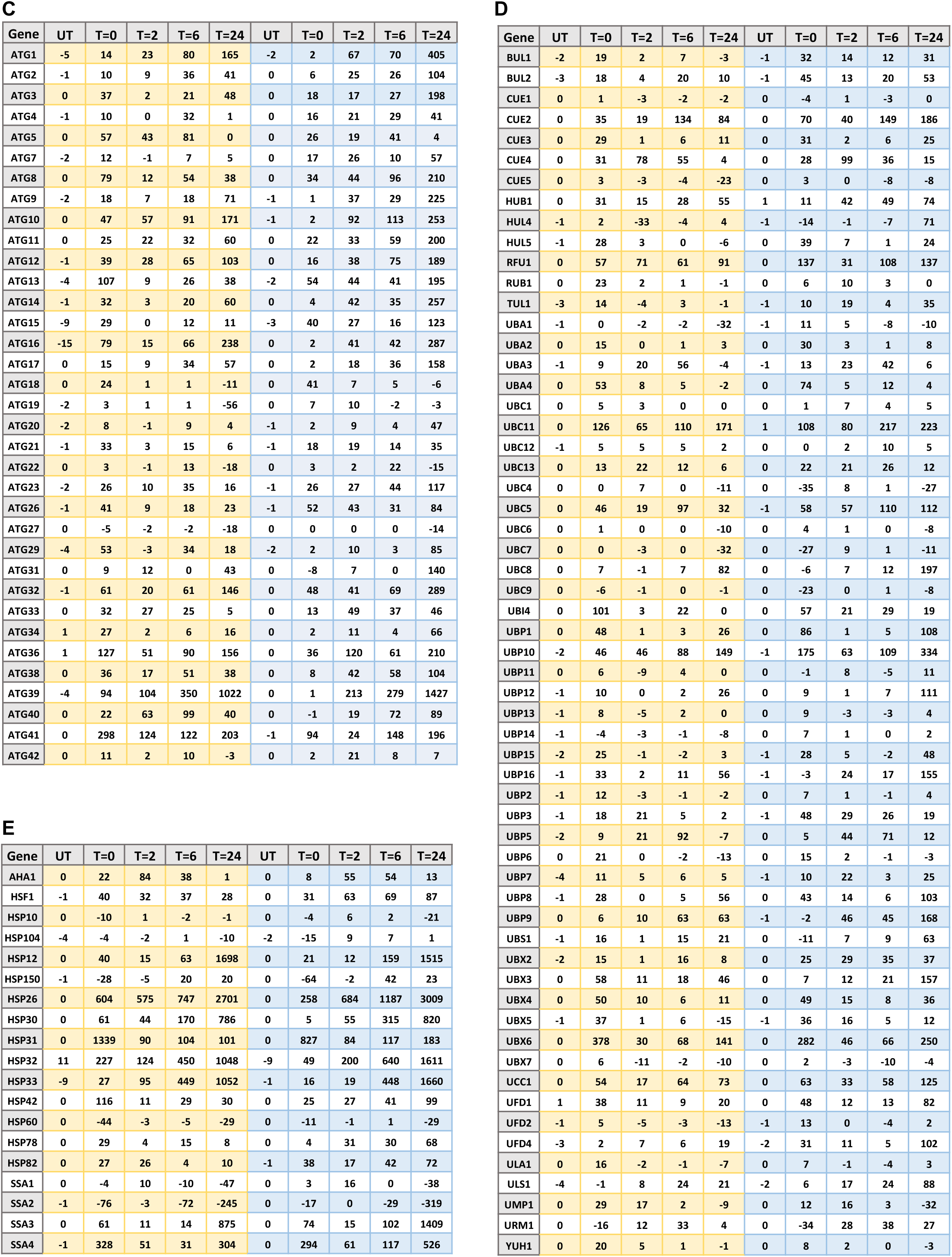
Transcriptome analysis of WT and *mgt1Δ* yeast cells. **A.** Heat shock genes are upregulated at the transcript level in cells treated with MNNG. **B.** Genes related to ubiquitin-mediated protein degradation are upregulated at the transcript level in cells treated with MNNG as well. In both cases, transcripts remain upregulated at higher levels in cells that cannot repair O^6^-MeG lesions. **C-E.** Average difference in gene expression between untreated samples (UT) and samples treated with MNNG at various timepoints. T=0 was taken immediately after exposure, T=2 was taken 2 hours after exposure, T=6 was taken 6 hours after exposure, and T=24 was taken 24 hours after exposure. WT cells are depicted in yellow, and MGT1Δ cells are depicted in blue. **C** depicts autophagy genes, **D** Ubiquitin related genes and **E** heat shock genes.

**Figure S12.**
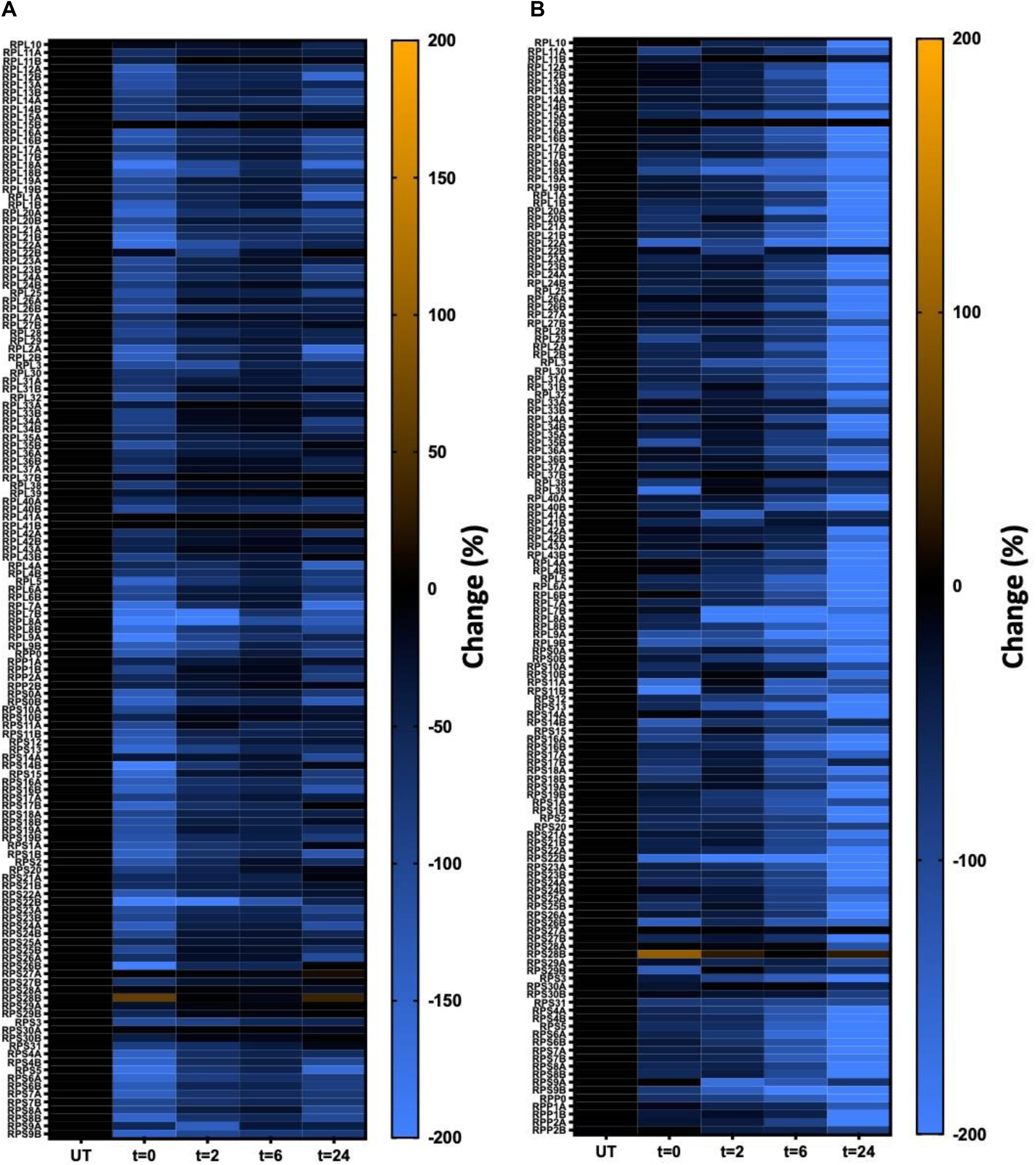
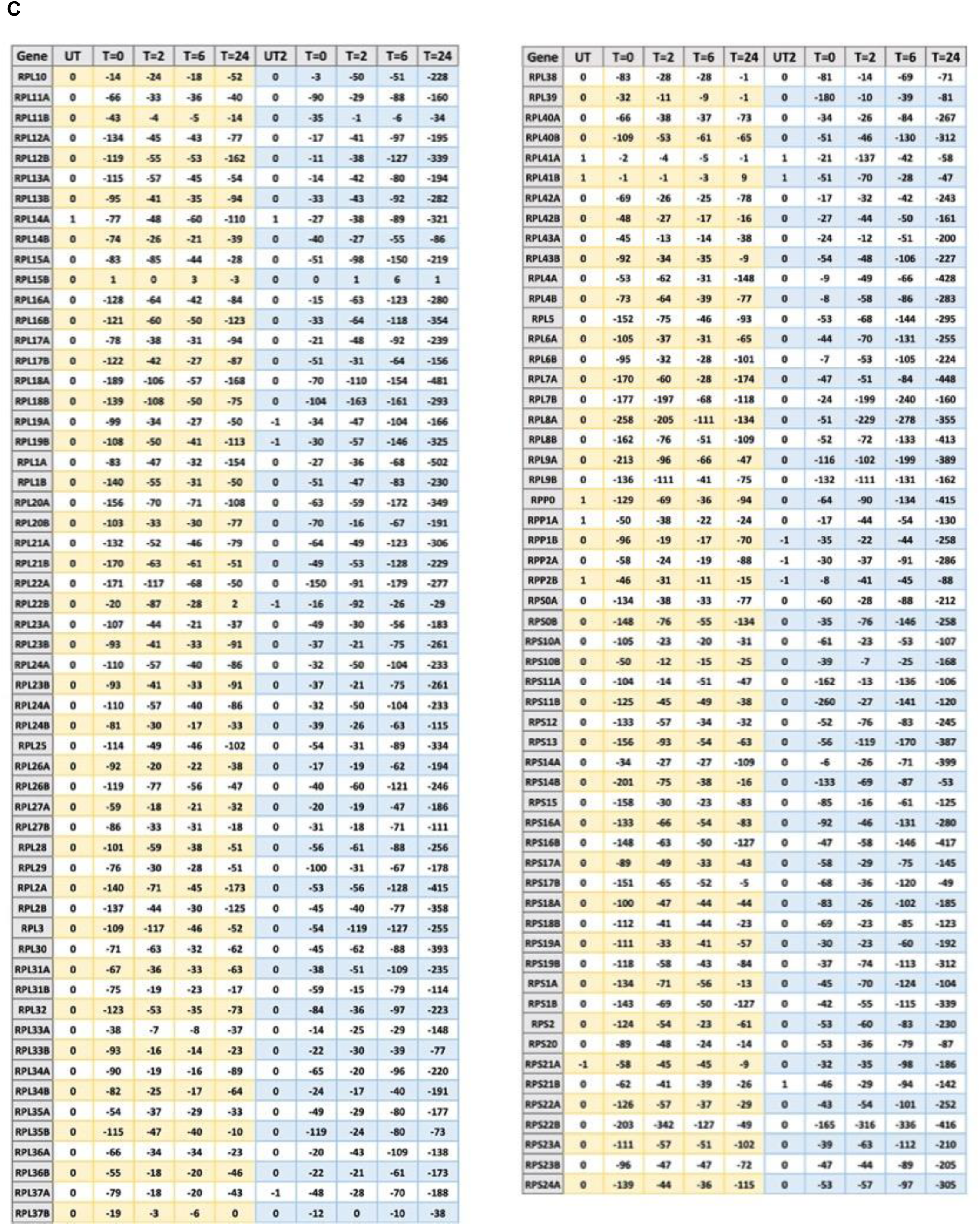
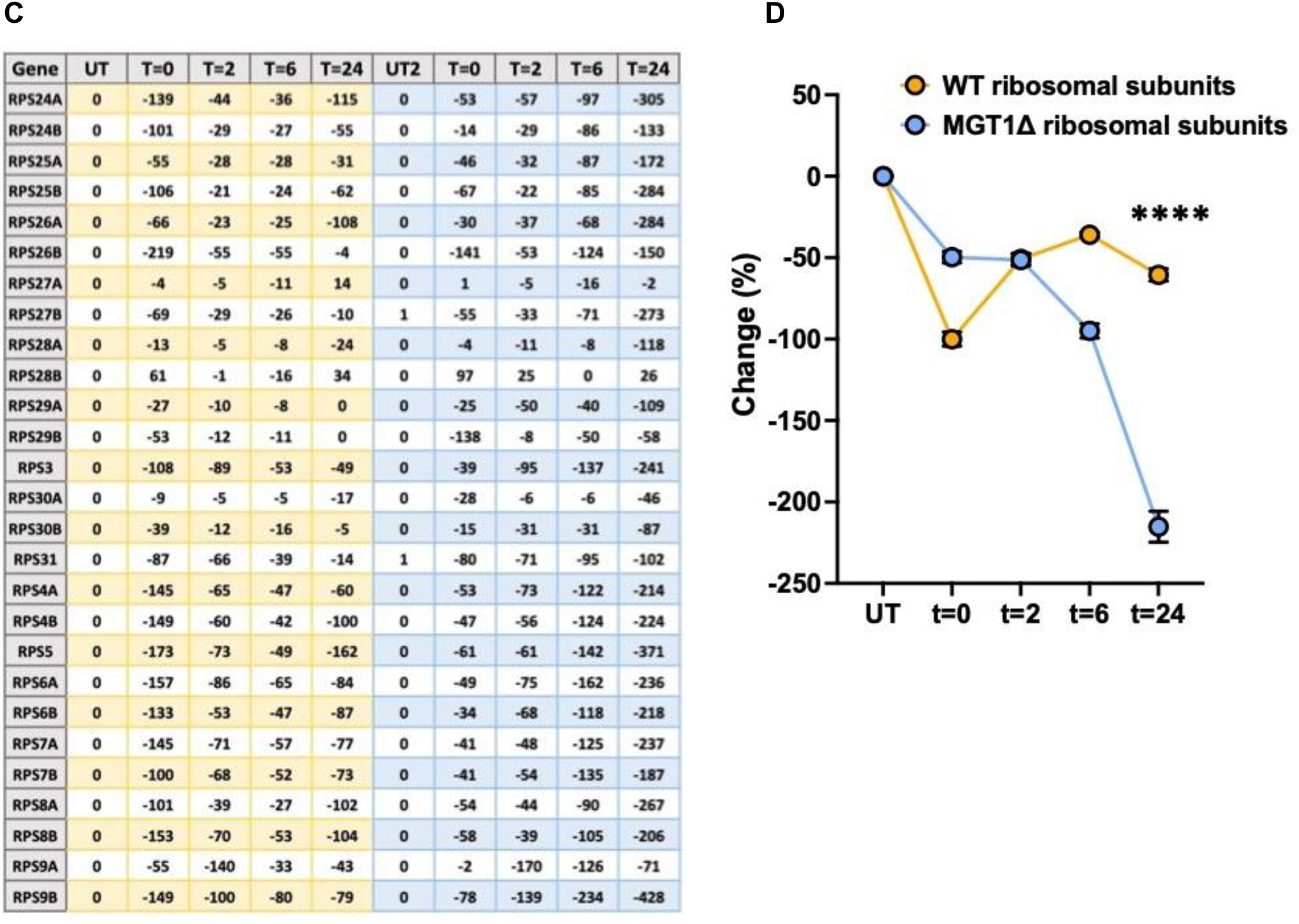
Transcriptome analysis of WT and *mgt1Δ* cells. **A.** Ribosomal subunits are downregulated at the transcript level in WT cells treated with MNNG. **B.** Ribosomal subunits genes are also downregulated at the transcript level in *mgt1Δ* cells treated with MNNG, which display an extended period of downregulation compared to WT cells. **C.** Average difference in gene expression between untreated samples (UT) and samples treated with MNNG at various timepoints. T=0 was taken immediately after exposure, T=2 was taken 2 hours after exposure, T=6 was taken 6 hours after exposure, and T=24 was taken 24 hours after exposure. WT cells are depicted in yellow, and MGT1Δ cells are depicted in blue **D.** Average percentage decrease of all ribosomal subunits depicted in **A-C**. **** = P<0.0001 according to a paired t-test. Error bars indicate standard error of the mean.

**Figure S13.**
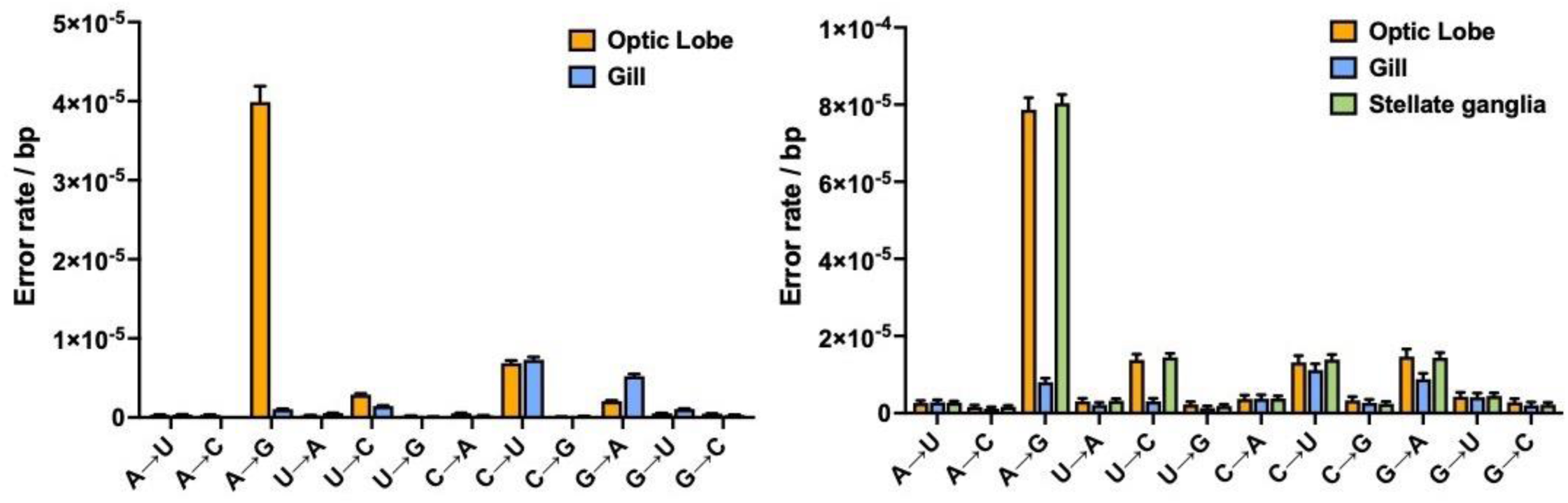
A. Error rate and spectra of squid and octopi. **A.** The optic lobe of squids (n=1) display a substantial increase in A to G substitutions caused by off-target A to I editing by high ADAR1 expression. ADAR1 expression is low in the gills (n=1) and as a result little off-target editing is seen. **B.** The optic lobe and stellate ganglia of octopi (which express ADAR1 at high levels) display substantial A to I editing (n=1), but the gills (which express ADAR1 at low levels) do not (n=1). Error bars indicate upper and lower limits.

## METHODS

### H1 hESC Cell culture

H1 hESCs were purchased from WiCell in Wisconsin (WA01) and cultured in TeSR medium in Matrigel coated 10cm plates. Cells were grown at 5% O_2_ tension to better mimic the conditions inside the human body and reduce oxidative damage as a result of normoxic conditions. To passage cells and prior to collection of RNA and DNA, cells were gently treated with 2 µg/ml Dispase mixed with DMEM/F12, washed with PBS and scraped off the plate using a glass pipette. DNA and RNA were then isolated with standard phenol chloroform and Trizol methods.

### Library Construction and Sequencing

Library preparation 1100ng of enriched mRNA was fragmented with the NEBNext RNase III RNA Fragmentation Module (E6146S) for 25 minutes at 37°C. RNA fragments were then purified with an Oligo Clean & Concentrator kit (D4061) by Zymo Research according to the manufacturer’s recommendations, except that the columns were washed twice instead of once. The fragmented RNA was then circularized with RNA ligase 1 in 20 µl reactions (NEB, M0204S) for 2 hours at 25°C after which the circularized RNA was purified with the Oligo Clean & Concentrator kit (D4061) by Zymo Research. The circular RNA templates were then reverse transcribed in a rolling-circle reaction by first incubating the RNA with for 10 minutes at 25°C to allow the random hexamers used for priming to bind to the templates. Then, the reaction was shifted to 42°C for 20 minutes to allow for primer extension and cDNA synthesis. Second strand synthesis and the remaining steps for library preparation were then performed with the NEBNext Ultra RNA Library Prep Kit for Illumina (E7530L) and the NEBNext Multiplex Oligos for Illumina (E7335S, E7500S) according to the manufacturer’s protocols. Briefly, cDNA templates were purified with the Oligo Clean & Concentrator kit (D4061) by Zymo Research and incubated with the second strand synthesis kit from NEB (E6111S). Double-stranded DNA was then entered into the end-repair module of RNA Library Prep Kit for Illumina from NEB, and size selected for 500-700 bp inserts using AMPure XP beads. These molecules were then amplified with Q5 PCR enzyme using 11 cycles of PCR, using a two-step protocol with 65°C primer annealing and extension and 95°C melting steps. Sequencing data was converted to industry standard Fastq files using BCL2FASTQv1.8.4.

### Error Identification

We have developed a robust bioinformatics pipeline to analyze circ-seq datasets and identify transcription errors with high sensitivity(*12, 17*). First, tandem repeats are identified within each read (minimum repeat size: 30nt, minimum identity between repeats: 90%), and a consensus sequence of the repeat unit is built. Next, the position that corresponds to the 5′ end of the RNA template is identified (the RT reaction is randomly primed, so cDNA copies can “start” anywhere on the template) by searching for the longest continuous mapping region. The consensus sequence is then reorganized to start from the 5′ end of the original RNA fragment, mapped against the genome with tophat (version 2.1.0 with bowtie 2.1.0) and all non-perfect hits go through a refining algorithm to search for the location of the 5′ end before being mapped again. Finally, every mapped nucleotide is inspected and must pass **5** checks to be retained: **1)** it must be part of at least 3 repeats generated from the original RNA template; **2)** all repeats must make the same base call; **3)** the sum of all qualities scores of this base must be >100; **4)** it must be >2 nucleotides away from both ends of the consensus sequence; **5)** each base must be covered by ≥100 reads with <1% of these reads supporting a base call different from the reference genome. This final step filters out polymorphic sites and intentional potential RNA-editing events. For example, if a base call is different from the reference genome, but is present in 50 out of 100 reads, it is not labeled as an error but as a heterozygous mutation. A similar rationale applies to low-level mutations and RNA editing events. These thresholds were altered to detect different types of editing events, including common editing events. Each read containing ≥1 mismatch is filtered through a second refining and mapping algorithm to ensure that errors in calling the position of the 5′ end cannot contribute to false positives. The error rate is then calculated as the number of mismatches divided by the total number of bases that passed all quality thresholds.

### Brain organoid culture and generation

H1 ESC colonies were maintained with daily media change in mTeSR (STEMCELL Technologies, #85850), supplemented with a final concentration of 5 μM XAV-939 (STEMCELL Technologies, #72672) on 1:100 geltrex (GIBCO, #A1413301) coated tissue culture plates (CELLTREAT, #229106) and passaged using ReLeSR (STEMCELL Technologies, #100-0484). Cells were maintained below passage 50 and periodically karyotyped via the G-banding Karyotype Service at Children’s Hospital Los Angeles. To generate dorsally patterned forebrain organoids, we modified the method previously described in Kadoshima *et al* (*68*). We eliminated the need for growth under 40% O2, the need for cell aggregates to be periodically bisected, and the use of high O_2_ penetration dishes, by adapting the cultures to growth in spinner-flask bioreactors. Specifically, on day 0, feeder-free cultured human PSCs, 80–90% confluent, were dissociated to single cells with Accutase (Gibco), and 9,000 cells per well were reaggregated in ultra-low cell-adhesion 96-well plates with V-bottomed conical wells (sBio PrimeSurface plate; Sumitomo Bakelite) in Cortical Differentiation Medium (CDM) I, containing Glasgow-MEM (Gibco), 20% Knockout Serum Replacement (Gibco), 0.1 mM Minimum Essential Medium non-essential amino acids (MEM-NEAA) (Gibco), 1 mM pyruvate (Gibco), 0.1 mM 2-mercaptoethanol (Gibco), 100 U/mL penicillin, and 100 μg/mL streptomycin (Corning). From day 0 to day 6, ROCK inhibitor Y-27632 (Millipore) was added to the medium at a final concentration of 20 μM. From day 0 to day 18, Wnt inhibitor IWR1 (Calbiochem) and TGFβ inhibitor SB431542 (Stem Cell Technologies) were added at a concentration of 3 μM and 5 μM, respectively. From day 18, the floating aggregates were cultured in ultra-low attachment culture dishes (Corning) under orbital agitation (70 rpm) in CDM II, containing DMEM/F12 medium (Gibco), 2mM Glutamax (Gibco), 1% N2 (Gibco), 1% Chemically Defined Lipid Concentrate (Gibco), 0.25 μg/mL fungizone (Gibco), 100 U/mL penicillin, and 100 μg/mL streptomycin. On day 35, cell aggregates were transferred to spinner-flask bioreactors (Corning) and maintained at 56 rpm, in CDM III, consisting of CDM II supplemented with 10% fetal bovine serum (FBS) (GE-Healthcare), 5 μg/mL heparin (Sigma), and 1% Matrigel (Corning). From day 70, organoids were cultured in CDM IV, consisting of CDM III supplemented with B27 supplement (Gibco) and 2% Matrigel.

### Neuronal culture and generation

H1ESCs were grown to confluency, split with accutase and seeded at a density of 3×10^5^ cells per well of a coated 6-well plate in mTeSR supplemented with 10μM rock inhibition. Cells were then transduced with hNGN2 and RTTA lentiviruses to obtain >90% infection efficiency using 4 μg/ml polybrene. mTeSR was changed daily until the cells were ready to split into a single-cell suspension with accutase, and the seeded directly into N2 media, so that approximately 1.2 × 10^6^ cells were present per 10cm dish. After 1 day, the N2 media was replaced with N2 media supplemented with puromycin at a concentration of 0.7 ug/ml to enable selection for transduced clones, which is complemented 2 days later with B27. The media was then replaced with N2 B27 media supplemented with 2 uM Ara-C (1-β-D-Arabino-furanosylcytosine) with ½ media change every other day. Cells were then allowed to grow and mature into neurons for 2 weeks before RNA isolation and error measurements.

#### Lentiviral generation and transduction

HEK293T cells were plated at 25% confluency and then transfected with plasmids that carry WT of or mutant versions of various proteins using Origene’s lentiviral packaging kit (TR30037). Medium was replaced after 18 hours of incubation and viral particles were harvested 24 and 48 hours later and filtered through a 0.45μm PES filter. The particles were then concentrated using a sucrose gradient in a Beckman ultracentrifuge at 70,000xg for 2.5 hours at 4°C. Afterwards, the viral pellets were resuspended in 25μL ice cold dPBS for every 15mL of viral medium spun down. AG10215 fibroblasts and U87 glioblastoma cells were then transduced with the concentrated viral particles at various MOIs 5 in antibiotic-free complete medium with 8μg/mL of polybrene. Cells were incubated for 18-24 hours before medium was changed to complete medium. Antibiotic selection for transduced cells began 48 hours after transduction and fluorescence assessed with a Leica Stellaris confocal microscope.

#### Mouse neural stem cell culture

Cells were cultured at 37°C in 5% CO_2_ and 5% O_2_ on PLO- and laminin-coated wells in serum-free media (NSC media) containing 1x DMEM/F12 (Invitrogen, 10565018), 1x pen/strep (Invitrogen 15140122), 1xB27 (Invitrogen, 17504044), 20ng/ml FGF2 (PeproTech, 100-18B), 20ng/ml EGF (PeproTech, AF-100-15) and 5μg/ml heparin (Sigma, H3149). For quiescence induction, cells were grown for at least 3 days in the same medium as described above, but without FGF2 and with the addition of 50ng/ml BMP-4 (Fisher Scientific, 5020BP010).

#### MGMT protein levels

Immunoblotting: 20ug of nuclear lysates were boiled at 75°C under denatured conditions and resolved on 4-20% gradient gels. Proteins were electroblotted using a Criterion blotter (Bio-Rad Laboratories, Hercules, CA) and transferred onto 0.45um polyvinyl difluoride membranes. Membranes were stained using Revert 700 fluorescent protein stain as a loading control and imaged prior to blocking with Intercept blocking buffer (LI-COR Biosciences, Lincoln, NE). Membranes were incubated overnight for 16 hours with 1:500 MGMT primary antibody (67476-1-Ig; Proteintech, Rosemead, IL). Membranes incubated with IRDye 800CW and/or 700CW secondary antibodies and visualized with a LI-COR Odyssey C1920. Densitometry was quantified with ImageJ and normalized by total protein per lane.

#### Protein expression and purification

TP53 (aa 92-292) clones in Pet28a were transformed into Rosetta DE3 pLysS competent cells (Novagen) and induced by 1mM IPTG at 18 ° overnight. Then they were purified by Ni-NTA agarose (Qiagen). After additional purification by Mono S column (GE Healthcare) and buffer exchange, they were loaded onto Superdex 75 gel filtration column (GE Healthcare) running on an ÄKTA FPLC system. SOD1 (aa 1-154) clones in Pet28a were transformed into Rosetta DE3 pLysS competent cells (Novagen) and induced by 1mM IPTG at 18 degree overnight. Then they were purified by Ni-NTA agarose (Qiagen), and which were further purified by Superdex 75 gel filtration column (GE Healthcare) running on an ÄKTA FPLC system

#### Transmission electron microscopy, atomic force microscopy, fiber growth and hanging drop method

For TEM, protein samples were spotted on carbon-coated Formvar grid (Ted Pella). The samples were stained with nanoW/uranyl acetate before air drying. The images were taken on Talos F200C G2 at 80 kV at the Core Center of Excellence in Nano Imaging (CNI). For AFM, protein samples of different seeding conditions were spotted on MICA sheets before loading on Dimension Icon(Bruker), with SCANASYST-AIR probe in ScanAsyst mode. Different dilution ratios were tested for the best visualization condition. To monitor fiber growth inside wells or by the hanging drop method, protein samples of different seeding and dilution conditions were set up either in wells or hanging drop manner. All samples were observed under polarized light to ensure fiber structure existence. A range of high concentrations of NaCl was used in the mother liquor of the hanging drop tray to induce the necessary evaporation.

### Single cell experiments

Cells were treated for 1 hour (mNSCs) or 40 minutes (yeast) with 10μg/ml MNNG. Cells were then counted with a MacsQuant cell counter and loaded onto 10xGenomics chip for GEM preparation according to 10x Genomics protocols, so that approximately 5,000 cells would be captured inside GEMs. expected for. For yeast cells, 8,000 cells were loaded with the expectation that that would result in 5,000 successful GEMs as well. In addition, 1μl of zymolyase was added to the cell suspension to facilitate the removal of the yeast cell wall. Results were then analyzed by CellRanger software, and on average, 2,000-9,000 single cells passed QC thresholds and were successfully sequenced for each replicate.

### Seurat processing for mNSC single cell RNA-seq

CellRanger output folders were imported for processing in R v3.6.3 using Seurat v3.2.2(*69*). Runs from 2 independent batches were merged together for analysis. To retain only high-quality cells, we applied filters nFeature_RNA > 1000 & percent.mito < 20. To determine likely cell cycle stage, a list of mouse cell cycle genes was obtained from the Seurat Vignettes (https://www.dropbox.com/s/3dby3bjsaf5arrw/cell_cycle_vignette_files.zip?dl=1), derived from a mouse study(*70*). Cell cycle phase was predicted using these genes, and using the function CellCycleSorting to assign cell cycle scores to each cell. Likely Doublets were identified using DoubletFinder 2.0(*71*), and removed from downstream processing. Reciprocal PCA was used to integrate data from the 2 cohortsand mitigate batch effects, using the top 7500 most variable genes and with k = 10. To determine whether proteostasis-related terms were differentially regulated in response to DNA-damage in quiescent NSCs at the single-cell level, we leveraged the UCell robust single-cell gene signature scoring metric implemented through R package ‘UCell’ 1.3.1(*72*). Cell-wise UCell scores were computed for selected GO terms related to proteostasis. Genes associated to these GO terms were obtained from ENSEMBL Biomart (version 109; accessed 2023-04-22), to retain relationships with all evidence codes except NAS/TAS, which nonexistent experimental support. For analysis of statistical significance, we used ANOVA to compare the distribution of UCell scores across time points and is reported for each gene set, and p-values were corrected for multiple hypothesis testing using the Benjamini-Hochberg method.

#### Pseudo-allele detection

Sequencing reads are first processed with the Cell Ranger Pipeline100. For each cell, reads with the same UMI (*i.e.* PCR duplicates) are collapsed into a consensus sequence, which is incorporated into a pileup file summarizing the sequence of each unique transcript at each genomic position in each cell. Positions covered by at least 40 unique transcripts are retained for downstream analysis and those with at least 10% of unique bases divergent from genomic DNA are compiled into a final output file for each cell.

## REFERENCES

1. C. Lopez-Otin, M. A. Blasco, L. Partridge, M. Serrano, G. Kroemer, The hallmarks of aging. Cell 153, 1194–1217 (2013).

2. M. S. Hipp, P. Kasturi, F. U. Hartl, The proteostasis network and its decline in ageing. Nat Rev Mol Cell Biol 20, 421–435 (2019).

3. D. Eisenberg, M. Jucker, The amyloid state of proteins in human diseases. Cell 148, 1188–1203 (2012).

4. C. Scheckel, A. Aguzzi, Prions, prionoids and protein misfolding disorders. Nature reviews. Genetics 19, 405–418 (2018).

5. G. A. P. de Oliveira et al., The Status of p53 Oligomeric and Aggregation States in Cancer. Biomolecules 10, (2020).

6. M. A. Gertz, A. Dispenzieri, T. Sher, Pathophysiology and treatment of cardiac amyloidosis. Nat Rev Cardiol 12, 91–102 (2015).

7. K. L. Moreau, J. A. King, Protein misfolding and aggregation in cataract disease and prospects for prevention. Trends Mol Med 18, 273–282 (2012).

8. L. M. Dember, Amyloidosis-associated kidney disease. J Am Soc Nephrol 17, 3458–3471 (2006).

9. R. Schroder, Protein aggregate myopathies: the many faces of an expanding disease group. Acta Neuropathol 125, 1–2 (2013).

10. N. Gregersen, P. Bross, S. Vang, J. H. Christensen, Protein misfolding and human disease. Annu Rev Genomics Hum Genet 7, 103–124 (2006).

11. E. Garcion, B. Wallace, L. Pelletier, D. Wion, RNA mutagenesis and sporadic prion diseases. J Theor Biol 230, 271–274 (2004).

12. J. F. Gout et al., The landscape of transcription errors in eukaryotic cells. Sci Adv 3, e1701484 (2017).

13. M. Vermulst et al., Transcription errors induce proteotoxic stress and shorten cellular lifespan. Nat Commun 6, 8065 (2015).

14. T. T. Saxowsky, P. W. Doetsch, RNA polymerase encounters with DNA damage: transcription-coupled repair or transcriptional mutagenesis? Chemical reviews 106, 474–488 (2006).

15. S. B. Prusiner, Scrapie prions. Annu Rev Microbiol 43, 345–374 (1989).

16. J. Brettschneider, K. Del Tredici, V. M. Lee, J. Q. Trojanowski, Spreading of pathology in neurodegenerative diseases: a focus on human studies. Nat Rev Neurosci 16, 109–120 (2015).

17. C. Fritsch, J. P. Gout, M. Vermulst, Genome-wide Surveillance of Transcription Errors in Eukaryotic Organisms. Journal of visualized experiments : JoVE, (2018).

18. A. Acevedo, R. Andino, Library preparation for highly accurate population sequencing of RNA viruses. Nature protocols 9, 1760–1769 (2014).

19. C. Chung et al., The fidelity of transcription in human cells. Proc Natl Acad Sci U S A 120, e2210038120 (2023).

20. S. Mead, S. Lloyd, J. Collinge, Genetic Factors in Mammalian Prion Diseases. Annu Rev Genet 53, 117–147 (2019).

21. C. Van Cauwenberghe, C. Van Broeckhoven, K. Sleegers, The genetic landscape of Alzheimer disease: clinical implications and perspectives. Genet Med 18, 421–430 (2016).

22. H. P. Nguyen, C. Van Broeckhoven, J. van der Zee, ALS Genes in the Genomic Era and their Implications for FTD. Trends Genet 34, 404–423 (2018).

23. M. J. Landrum et al., ClinVar: improving access to variant interpretations and supporting evidence. Nucleic Acids Res 46, D1062–D1067 (2018).

24. P. D. Stenson et al., The Human Gene Mutation Database: towards a comprehensive repository of inherited mutation data for medical research, genetic diagnosis and next-generation sequencing studies. Hum Genet 136, 665–677 (2017).

25. T. Sato et al., Identification of two novel mutations in the Cu/Zn superoxide dismutase gene with familial amyotrophic lateral sclerosis: mass spectrometric and genomic analyses. J Neurol Sci 218, 79–83 (2004).

26. T. J. Kwiatkowski, Jr., et al., Mutations in the FUS/TLS gene on chromosome 16 cause familial amyotrophic lateral sclerosis. Science 323, 1205–1208 (2009).

27. D. B. Rowe et al., Novel prion protein gene mutation presenting with subacute PSP-like syndrome. Neurology 68, 868–870 (2007).

28. M. O. Kim, L. T. Takada, K. Wong, S. A. Forner, M. D. Geschwind, Genetic PrP Prion Diseases. Cold Spring Harb Perspect Biol 10, (2018).

29. H. An et al., ALS-linked FUS mutations confer loss and gain of function in the nucleus by promoting excessive formation of dysfunctional paraspeckles. Acta Neuropathol Commun 7, 7 (2019).

30. H. Wang et al., Mutant FUS causes DNA ligation defects to inhibit oxidative damage repair in Amyotrophic Lateral Sclerosis. Nat Commun 9, 3683 (2018).

31. S. Ishigaki, G. Sobue, Importance of Functional Loss of FUS in FTLD/ALS. Front Mol Biosci 5, 44 (2018).

32. C. F. Sephton et al., Activity-dependent FUS dysregulation disrupts synaptic homeostasis. Proc Natl Acad Sci U S A 111, E4769–4778 (2014).

33. S. L. Rulten et al., PARP-1 dependent recruitment of the amyotrophic lateral sclerosis-associated protein FUS/TLS to sites of oxidative DNA damage. Nucleic Acids Res 42, 307–314 (2014).

34. E. Srinivasan, R. Rajasekaran, A Systematic and Comprehensive Review on Disease-Causing Genes in Amyotrophic Lateral Sclerosis. J Mol Neurosci 70, 1742–1770 (2020).

35. S. Parakh, J. D. Atkin, Protein folding alterations in amyotrophic lateral sclerosis. Brain Res 1648, 633–649 (2016).

36. M. Lynch et al., Genetic drift, selection and the evolution of the mutation rate. Nature reviews. Genetics 17, 704–714 (2016).

37. J. A. Toombs et al., De novo design of synthetic prion domains. Proc Natl Acad Sci U S A 109, 6519–6524 (2012).

38. I. Le Ber et al., hnRNPA2B1 and hnRNPA1 mutations are rare in patients with “multisystem proteinopathy” and frontotemporal lobar degeneration phenotypes. Neurobiol Aging 35, 934 e935–936 (2014).

39. H. J. Kim et al., Mutations in prion-like domains in hnRNPA2B1 and hnRNPA1 cause multisystem proteinopathy and ALS. Nature 495, 467–473 (2013).

40. V. Espinosa Angarica et al., PrionScan: an online database of predicted prion domains in complete proteomes. BMC Genomics 15, 102 (2014).

41. A. K. Lancaster, A. Nutter-Upham, S. Lindquist, O. D. King, PLAAC: a web and command-line application to identify proteins with prion-like amino acid composition. Bioinformatics 30, 2501–2502 (2014).

42. S. Pawlicki, A. Le Bechec, C. Delamarche, AMYPdb: a database dedicated to amyloid precursor proteins. BMC Bioinformatics 9, 273 (2008).

43. T. Eneqvist, K. Andersson, A. Olofsson, E. Lundgren, A. E. Sauer-Eriksson, The beta-slip: a novel concept in transthyretin amyloidosis. Molecular cell 6, 1207–1218 (2000).

44. H. R. Fryer, A. R. McLean, There is no safe dose of prions. PLoS One 6, e23664 (2011).

45. T. T. Saxowsky, K. L. Meadows, A. Klungland, P. W. Doetsch, 8-Oxoguanine-mediated transcriptional mutagenesis causes Ras activation in mammalian cells. Proc Natl Acad Sci U S A 105, 18877–18882 (2008).

46. D. Bregeon, Z. A. Doddridge, H. J. You, B. Weiss, P. W. Doetsch, Transcriptional mutagenesis induced by uracil and 8-oxoguanine in Escherichia coli. Molecular cell 12, 959–970 (2003).

47. A. Viswanathan, H. J. You, P. W. Doetsch, Phenotypic change caused by transcriptional bypass of uracil in nondividing cells. Science 284, 159–162 (1999).

48. M. D. Wyatt, D. L. Pittman, Methylating agents and DNA repair responses: Methylated bases and sources of strand breaks. Chem Res Toxicol 19, 1580–1594 (2006).

49. Y. Mu, F. H. Gage, Adult hippocampal neurogenesis and its role in Alzheimer’s disease. Mol Neurodegener 6, 85 (2011).

50. S. Teuber-Hanselmann, K. Worm, N. Macha, A. Junker, MGMT-Methylation in Non-Neoplastic Diseases of the Central Nervous System. Int J Mol Sci 22, (2021).

51. S. L. Gerson, MGMT: its role in cancer aetiology and cancer therapeutics. Nat Rev Cancer 4, 296–307 (2004).

52. J. Chung et al., Genome-wide association and multi-omics studies identify MGMT as a novel risk gene for Alzheimer’s disease among women. Alzheimers Dement, (2022).

53. C. Fritsch et al., Genome-wide surveillance of transcription errors in response to genotoxic stress. Proc Natl Acad Sci U S A 118, (2021).

54. C. Chung et al., Evolutionary conservation of the fidelity of transcription. Nat Commun 14, 1547 (2023).

55. A. Konopka, J. D. Atkin, DNA Damage, Defective DNA Repair, and Neurodegeneration in Amyotrophic Lateral Sclerosis. Front Aging Neurosci 14, 786420 (2022).

56. N. Liscovitch-Brauer et al., Trade-off between Transcriptome Plasticity and Genome Evolution in Cephalopods. Cell 169, 191–202 e111 (2017).

57. J. Tao, D. E. Bauer, R. Chiarle, Assessing and advancing the safety of CRISPR-Cas tools: from DNA to RNA editing. Nat Commun 14, 212 (2023).

58. B. Falcon et al., Tau filaments from multiple cases of sporadic and inherited Alzheimer’s disease adopt a common fold. Acta Neuropathol 136, 699–708 (2018).

59. C. M. Tanner et al., Rotenone, paraquat, and Parkinson’s disease. Environ Health Perspect 119, 866–872 (2011).

60. A. Spivey, Rotenone and paraquat linked to Parkinson’s disease: human exposure study supports years of animal studies. Environ Health Perspect 119, A259 (2011).

61. P. S. Spencer, V. S. Palmer, G. E. Kisby, Western Pacific ALS-PDC: Evidence implicating cycad genotoxins. J Neurol Sci 419, 117185 (2020).

62. B. M. Verheijen, T. Hashimoto, K. Oyanagi, F. W. van Leeuwen, Deposition of mutant ubiquitin in parkinsonism-dementia complex of Guam. Acta Neuropathol Commun 5, 82 (2017).

63. R. M. Garruto, R. Yanagihara, D. C. Gajdusek, Disappearance of high-incidence amyotrophic lateral sclerosis and parkinsonism-dementia on Guam. Neurology 35, 193–198 (1985).

64. L. A. Farrer et al., Effects of age, sex, and ethnicity on the association between apolipoprotein E genotype and Alzheimer disease. A meta-analysis. APOE and Alzheimer Disease Meta Analysis Consortium. JAMA 278, 1349–1356 (1997).

65. F. W. van Leeuwen et al., Frameshift mutants of beta amyloid precursor protein and ubiquitin-B in Alzheimer’s and Down patients. Science 279, 242–247 (1998).

66. F. W. van Leeuwen, J. P. Burbach, E. M. Hol, Mutations in RNA: a first example of molecular misreading in Alzheimer’s disease. Trends Neurosci 21, 331–335 (1998).

67. H. H. Guo, J. Choe, L. A. Loeb, Protein tolerance to random amino acid change. Proc Natl Acad Sci U S A 101, 9205–9210 (2004).

68. T. Kadoshima et al., Self-organization of axial polarity, inside-out layer pattern, and species-specific progenitor dynamics in human ES cell-derived neocortex. Proc Natl Acad Sci U S A 110, 20284–20289 (2013).

69. A. Butler, P. Hoffman, P. Smibert, E. Papalexi, R. Satija, Integrating single-cell transcriptomic data across different conditions, technologies, and species. Nature biotechnology 36, 411–420 (2018).

70. M. S. Kowalczyk et al., Single-cell RNA-seq reveals changes in cell cycle and differentiation programs upon aging of hematopoietic stem cells. Genome Res 25, 1860–1872 (2015).

71. C. S. McGinnis, L. M. Murrow, Z. J. Gartner, DoubletFinder: Doublet Detection in Single-Cell RNA Sequencing Data Using Artificial Nearest Neighbors. Cell Syst 8, 329–337 e324 (2019).

72. M. Andreatta, S. J. Carmona, UCell: Robust and scalable single-cell gene signature scoring. Comput Struct Biotechnol J 19, 3796–3798 (2021).

